# Propagation Mapping: A Framework for Modeling Whole-Brain Propagation Patterns of Task-Evoked Activity

**DOI:** 10.1101/2025.09.27.678975

**Authors:** Jules R. Dugré

**Author notes:** **Corresponding authors:** Jules Roger Dugré, PhD.

## Abstract

Human brain mapping has traditionally relied on univariate approaches to characterize regional activity, whereas more recent work focuses on interactions between regions to capture network-level organization. Despite their parallel development, growing evidence suggests that integrating both approaches is critical for a comprehensive understanding of task-evoked brain activity. The present study introduces propagation mapping, an extension of activity flow mapping that model’s task-evoked brain activity as the propagation of regional signal amplitudes along whole-brain topological routes. This study aims to evaluate propagation maps as reliable neurobiological features for neuroimaging research. Using functional connectomes and structural covariance network derived from a large normative sample (n=1,000), propagation patterns of task-evoked activity were accurately captured (average *R*^2^ = 0.947, MAE=0.155, and RMSE=0.229) across 94 participants. Mapping performance remained stable across different task contrasts, parcellation atlases, and signal intensity and spatial distance between regions. Similar performance was observed at both the subject and group levels in an independent sample using the amplitude of low-frequency oscillations during resting-state (n=189). Importantly, despite its reliance on normative connectomes which could homogenize subject-specific variance, propagation mapping instead redistributed individual variance along propagation routes (Cohen’s *d* = 0.10, p=0.17). As a biologically comprehensive representation of brain organization, propagation mapping offers a powerful and user-friendly alternative to traditional regional analyses and provides new avenues for discovery in neurological and psychiatric neuroimaging research.

## 1 Introduction

From the earliest days of human brain mapping, researchers have relied on statistical parametric maps to infer brain structure and function. With the advent of MRI, voxelwise maps derived from different modalities including metabolism, brain morphometry, and BOLD signal amplitude have become central to neuroimaging (Fox et al., 1986; Friston et al., 1995; Wright et al., 1995). Mass-univariate analyses were initially preferred over multivariate models because measured hemodynamic responses can differ across brain regions, even if the underlying neuronal activity is the same (Friston, Penny, Ashburner, Kiebel, & Nichols, 2006). By analyzing each voxel separately, mass-univariate approaches avoided misattributing regional variations in hemodynamic signals to neuronal differences. Despite methodological advances, mass-univariate approaches for capturing regional BOLD activity have increasingly been criticized from a conceptual standpoint. Indeed, the approach relies on the erroneous assumption that brain regions operate as isolated units (Noble, Curtiss, Pessoa, & Scheinost, 2024; Segal et al., 2025), a premise that is difficult to reconcile with the billions of highly interconnected neurons in the human brain (Goriely, 2025). This highlights the importance of conceptualizing the brain as a complex system of interconnected regions to advance scientific discovery (Noble et al., 2024). In turn, functional connectivity (FC), which quantifies the temporal synchrony between signal fluctuations of two regions, offers a promising alternative to traditional regional approaches. Yet, this approach overlooks contributions of local neural activity that are critical for understanding brain function. Taken together, although commonly described as independent measures, activity and connectivity are increasingly recognized as complementary rather than mutually exclusive.

Since the early days of MRI (Biswal, Yetkin, Haughton, & Hyde, 1995), the amplitude of the BOLD signal has been shown to closely align with regional connectivity strength, which measures how functionally connected a given region is to the rest of the brain (Di et al., 2013; Garrett, Epp, Perry, & Lindenberger, 2018; Liang, Zou, He, & Yang, 2013; Sheng et al., 2021). This may be partly explained by the fact that brain regions with stronger FC to the rest of the brain have higher metabolic demands (e.g., oxygen, glucose) (Castrillon et al., 2023; Wu et al., 2009), which in turn reflect greater blood flow fluctuations. These findings highlight the need for approaches that integrate both regional activity and connectivity to provide a more comprehensive account of brain function. According to recent findings, task-evoked activity in a given brain region can be predicted by modeling how neural activity propagates from other regions (Cole, Ito, Bassett, & Schultz, 2016). Specifically, activity flow mapping estimates brain activity as the sum of interactions between the amplitudes of all other regions and their resting-state FC to the target region (Cocuzza, Sanchez-Romero, & Cole, 2022; Cole et al., 2016). This method has been extensively validated in healthy participants (Cocuzza et al., 2024; Cole et al., 2016; Cole, Ito, Cocuzza, & Sanchez-Romero, 2021; Sanchez-Romero, Ito, Mill, Hanson, & Cole, 2023), and its utility has also been demonstrated in psychiatric (Hearne et al., 2021) and neurological disorders (Cabalo et al., 2025). For instance, findings suggest that patients with epilepsy showed significant reductions in predicting brain activity during episodic and semantic memory tasks compared with healthy controls (Δr = 0.14–0.30, differences in correlation between observed and predicted activity) (Cabalo et al., 2025). Overall, these findings underscore the tight coupling between regional activity and connectivity and provide a novel framework for studying neurological and psychiatric disorders.

### 1.1 The Current Study

The goal of this paper is to introduce and validate a framework, as an extension of activity flow mapping, designed to model the propagation of task-evoked brain activity throughout the whole-brain. While activity flow mapping typically focuses on predicting task-evoked activity (Cocuzza et al., 2024; Cole et al., 2016; Cole et al., 2021; Sanchez-Romero et al., 2023), the propagation routes underlying these predictions remain largely unexplored. Indeed, previous studies have not examined whole-brain propagation patterns (i.e., region-to-region paths) as a potential neuroimaging feature for advancing scientific discovery. The intuition behind *propagation mapping* is that if task-evoked activity can be accurately predicted from a weighted sum function, then the weighted connectome used for prediction must accurately capture how task-evoked activity flows along whole-brain topological routes. Therefore, the present method aims to provide additional insights into how task-evoked activity propagates throughout the whole brain.

Although the spatial configuration of task-evoked activity varies widely across individuals, the underlying architecture appears largely stable (Cole, Bassett, Power, Braver, & Petersen, 2014; Gratton et al., 2018; Smith et al., 2009), serving as a blueprint for identifying individual differences in regional activity (Fox, 2018). Therefore, it was first hypothesized that FC estimated from a large normative sample may serve as a functional template to capture an individual’s propagation patterns of task-evoked regional activity. I further hypothesized that integrating structural covariance (SC) would help refine the propagation routes, yielding better mapping performance. Evidence indicates that brain morphometry co-varies across regions in ways that mirror FC (Alexander-Bloch, Giedd, & Bullmore, 2013; Mechelli, Friston, Frackowiak, & Price, 2005), suggesting that anatomical features such as volume may constrain and guide functional interactions.

The current manuscript is organized into two complementary studies to validate the method.

In Study 1, the aim was to identify the biological model that best captures how task-evoked activity flows throughout the whole brain. It was hypothesized that incorporating normative FC and SC estimates would serve as blueprints to identify propagation routes, thereby enhancing the mapping accuracy of subjects’ task-evoked activity. Performance was expected to remain stable across task contrasts, parcellation atlases, signal intensity, and spatial distance.

Study 2 aimed to evaluate the generalizability of propagation mapping in an independent sample using resting-state low-frequency oscillations. The first goal was to determine whether the propagation patterns observed at the group level remain consistent and robust. The second goal was to assess the impact of the use of normative connectomes to estimate propagation patterns on inter-individual variability. To test this, a method analogous to functional connectome fingerprinting (Amico & Goñi, 2018; Finn et al., 2015) was carried out to compare the identifiability of subjects based on their (1) observed amplitude vectors, and (2) estimated propagation maps. We hypothesized that propagation mapping would preserve both group-level structure and individual-specific variance.

## 2 Method

### 2.1 Propagation Mapping

#### 2.1.1 Normative Connectomes

One general hypothesis of the current study was that regional maps may operate on a general and stable functional architecture. Therefore, prediction of brain maps is assumed to benefit from using group-level connectomes derived from healthy subjects, as they show more stable and reliable FC estimates. Therefore, the prediction was conducted by utilizing group-level FC and SC estimates generated from 1000 healthy subjects (ages 18–35 years old, 50 % females) of the Brain Genomics Superstruct Project (Holmes et al., 2015; Yeo et al., 2011). Data acquisition and Information about preprocessing steps of functional neuroimaging data is available elsewhere (Yeo et al., 2011) and in Supplementary Material.

For each participant, the denoised brain activity signal was extracted from 426 regions, including 400 cortical (Schaefer et al., 2018), 14 subcortical (Fischl et al., 2002), 5 midbrain communities (Hansen et al., 2024) and 7 cerebellar (Buckner, Krienen, Castellanos, Diaz, & Yeo, 2011) regions for each participant. Correlations were then calculated between the activity of each region and all other regions, producing a connectivity matrix for each individual. These correlation values were converted into standardized scores (Fisher z-transformation) to allow comparison and subsequently averaged across all 1,000 participants, to generate a group-level FC matrix.

As detailed elsewhere (Dugré et al., 2025), structural data was preprocessed using the Computational Anatomy Toolbox (CAT12) within the Statistical Parametric Mapping 12 (SPM12) software (Gaser et al., 2024). As recommended (Gaser et al., 2024), weighted overall quality rating (IQR) and quartic mean z-scores were used to measure data homogeneity before and after pre-processing. Data with z-scores higher than 2 will be visually inspected to detect potential artefacts. For each subject, mean volume of each of the parcels from the 426 regions (see above) were extracted as well as total intracranial volume (TIV). Group-level structural covariance was estimated by conducting partial correlation between the mean volume of each of the 426 regions, while adjusting for TIV. Coefficients were converted to normally distributed z-scores using Fisher z-transformation.

Because the study aims to examine the stability of propagation mapping’s accuracy across different parcellation atlases, FC and SC matrices were generated for multiple widely used atlases: an anatomical atlas (Desikan-Kiliany Atlas, 68 cortical regions, (Desikan et al., 2006)), resting-state (Gordon Atlas, 333 parcels, (Gordon et al., 2016)) and multi-modal atlas (Glasser HCP-MMP1, 360 parcels, (Glasser et al., 2016)). Each parcellation was supplemented with the same 14 subcortical, 5 midbrain communities, and 7 cerebellar regions as described above to ensure comparability.

#### 2.1.2 Normative Connectomes

For each participant, propagation mapping was conducted to model the task-evoked propagation patterns of their brain activity. As shown in Figure 1, statistical images are first parcellated using a common parcellation atlas (e.g., Schaefer-400 7-Network). In a hypothetical model with equal contributions of FC and SC to propagation routes, the amplitude of region *i* is modeled as the sum of amplitudes from all other regions (*j* ≠ *i*), weighted by the average of their connections with region *i*. Specifically, the normative FC (*NormFC_ij_*), and normative SC (*NormSC_ij_*) strengths, derived from a large reference sample, contribute equally to defining the propagation routes of task-evoked activity. For both FC and SC, diagonals were set to zero to prevent data leakage. These connectivity-weighted terms are first averaged across modalities (functional and structural), and then symmetrized by averaging forward and backward terms to yield a single, undirected propagation value for each region pair that captures their total contribution. Building on previous work (Cole et al., 2016) SC was included as a potential biological constraint on FC, given its critical role in shaping brain function (Honey et al., 2009; Pang et al., 2023; Qing & Gong, 2016):

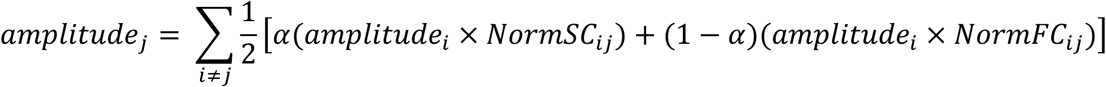

**Figure. 1.**
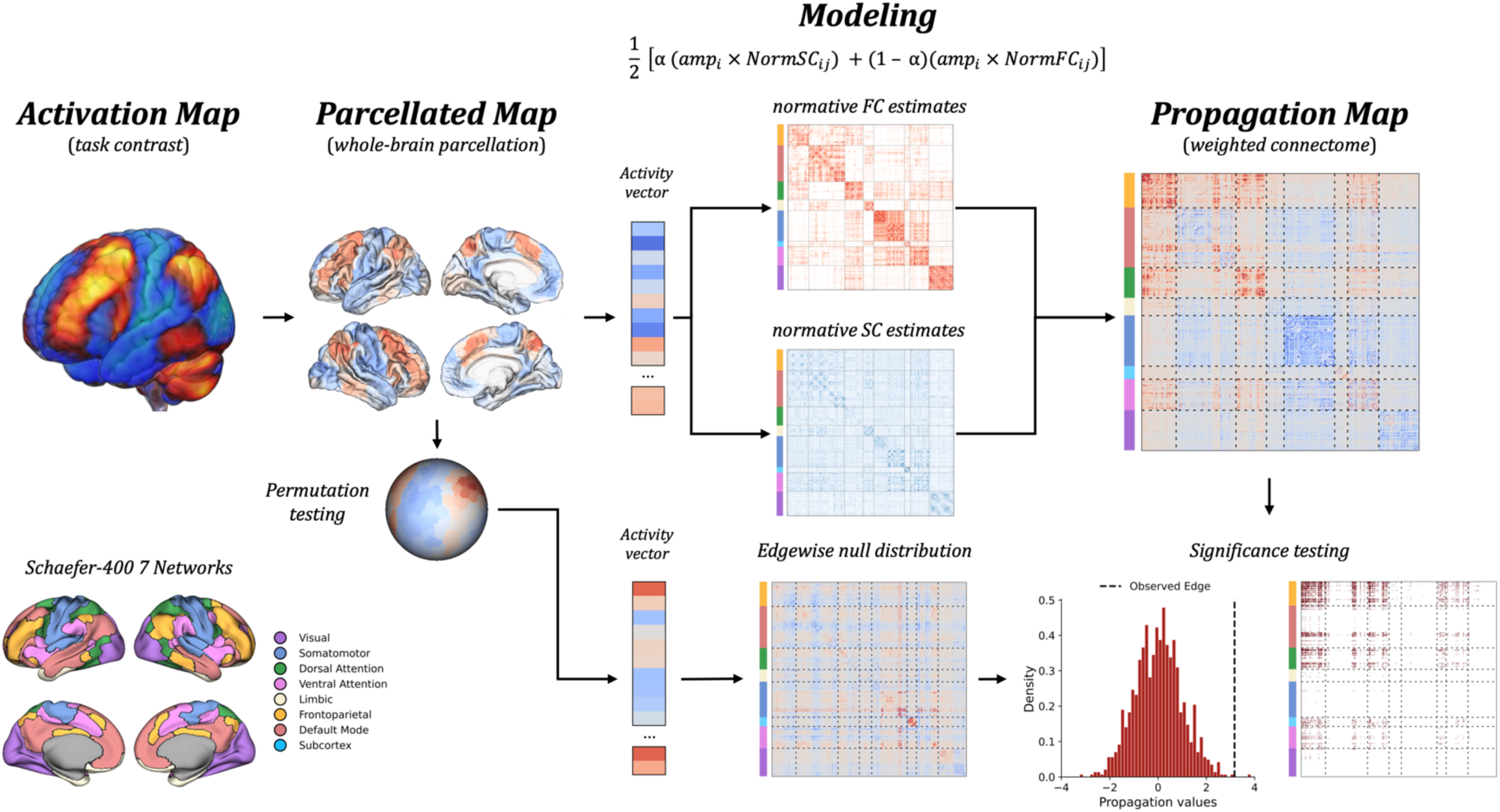
Methodological Overview of Propagation Mapping. This approach models the propagation patterns of task-evoked brain activity. First, the voxelwise statistical map is parcellated using commonly used atlases supplemented with 14 subcortical regions, 5 midbrain communities, and 7 cerebellar regions. The resulting regional activity vector serves as a multiplicative weighting term applied to connectivity-informed propagation routes. These routes are derived from functional connectivity (FC) and structural covariance (SC) matrices estimated from a large normative sample (n = 1000; GSP1000). The diagonals of these normative connectomes are set to zero to avoid self-influence. Averaging the weighted matrices yields a propagation map that reflects the subject-specific pattern of task-evoked signal propagation. To evaluate whether the approach captures connectivity-informed structure beyond regional proximity, spatial autocorrelation–preserving nulls were generated (1,000 permutations with Hungarian algorithm). Edgewise Z-scores of the observed propagation pattern were then computed relative to the combined null distributions, thresholded (p < 0.05), and binarized. Network colors in the connectivity matrices correspond to the seven networks defined by Schaefer et al., as shown in the bottom left. The Propagation Mapping Toolbox has been made freely available on Streamlit Cloud (https://propagation-mapping.streamlit.app/) or can be run locally (https://github.com/JulDugre/Propagation-Mapping).

To assess mapping performance, propagation values for each region *i* were summed across all other regions (*j* ≠ *i*) to generate a weighted degree (strength) feature vector reflecting the predicted task-evoked value of region *i*. For each participant, mapping accuracy was quantified using the coefficient of determination (*R*^2^), while mapping errors were quantified using mean absolute errors (MAE) and root mean squared error (RMSE) between the observed task-evoked activity pattern and the *predicted* map. A high mapping accuracy indicates that the weighted connectome, prior to the calculation of the weighted degree, effectively captures the propagation patterns underlying individual’s task-evoked activity pattern.

The Propagation Mapping Toolbox has been made freely available on Streamlit Cloud (https://propagation-mapping.streamlit.app/) or can be run locally (https://github.com/JulDugre/Propagation-Mapping).

### 2.2 Study 1

#### 2.2.1 Study sample and Design

A first study was conducted to test the reliability of propagation mapping across different subjects, task contrasts, parcellation atlases, and across short-range and long-range connections. For this initial study, data was obtained from the Brainomics/Localizer dataset (Papadopoulos Orfanos et al., 2017; Pinel et al., 2007), A subset of 94 participants was made openly available (see https://osf.io/vhtf6), which include 49 women and 45 men, with a mean age of 24.7 years old (18-49 years old). The Localizer study received ethical approval from their Local ethics committee, and all subjects provided informed consent (Papadopoulos Orfanos et al., 2017; Pinel et al., 2007). Data acquisition and preprocessing can be found elsewhere (Papadopoulos Orfanos et al., 2017; Pinel et al., 2007) and in Supplementary Material.

The Localizer task included ten trial types that were presented to participants. From these trial types, 6 task contrasts were selected in the current study given their neurobiological differences, to examine the predictive accuracy of propagation mapping: 1) Checkerboard (i.e., passive viewing of flashing checkerboards), 2) Sentence Reading (i.e., silently reading short visual sentences), 3) Sentence Listening (i.e., listening to short sentences). 4) Calculation (i.e., solving silently visual/auditory subtraction problems, 5) Left Button Press (i.e., pressing the left button with the left thumb according to the auditory/visual instruction), and 6) Right Button Press (i.e., pressing the right button with the right thumb according to the auditory/visual instruction).

#### 2.2.2 Statistical Analyses

##### Propagation Mapping Performance Across Biological Models

To identify the most optimal biological model for propagation mapping, I tested predictive accuracy across seven different models: (a) using the full FC matrix only; (b) using only positive edges of the FC matrix; (c) using the full SC matrix only; (d) using only positive edges of the SC matrix; (e) combining the full FC and SC matrices with equal weighting; (f) combining the positive edges of the FC and SC matrices with equal weighting; (g) combining the full FC and SC matrices with unequal weighting, defined by α, with all values of α ranging from 0 to 1 in increments of 0.02, to account for potential variations in the influence of SC on FC; and (e) combining only the positive edges of the FC and SC matrices using the same α weighting.

Accuracy of the biological models was assessed using *R*^2^, MAE, and RMSE. Critically, the stability of model accuracy was evaluated across six task contrasts (Checkerboard; Sentence Reading; Sentence Listening; Calculation; Left Button Press; Right Button Press), and across multiple parcellations. This was done to rule out the possibility that the findings were driven by a specific parcellation and to test that the method generalizes well across different parcellation atlases. To do so, 5 widely used cortical parcellations were included in the analyses including an anatomical (Desikan-Kiliany atlas, 68 cortical regions, (Desikan et al., 2006)), three resting-state (Gordon atlas, 333 parcels, (Gordon et al., 2016); Schaefer atlas, 400 parcels 7-Network, and 400 parcels 17-Network, (Schaefer et al., 2018)), and multi-modal (Glasser atlas, 360 parcels, (Glasser et al., 2016)) atlases. These were supplemented by additional 14 subcortical regions (Fischl et al., 2002), 5 midbrain communities (Hansen et al., 2024), and 7-cerebellar (Buckner et al., 2011) regions to allow comparability (i.e., 17 cerebellar regions for Schaefer 400 parcels 17 Network).

Once the most optimal biological model was identified for each parcellation, propagation mapping was repeated separately for regions within the lowest 25% and highest 25% of task-evoked signal intensities both in relative and absolute magnitude for each task contrast. This ensured that the mapping accuracy was not disproportionately driven by low-signal regions.

Moreover, to validate that the performance of the best biological model was not restricted to the current sample, 1,000 fMRI brain maps were randomly selected from NeuroVault, ensuring that they did not originate from the same collection to reduce the risk of bias.

Post hoc tests were conducted to examine whether FC and SC capture distinct and/or complementary neural features in identifying propagation routes. To this end, dominance analyses were performed to assess the relative contribution of three metrics to FC and SC: (a) Euclidean distance between the coordinates of the vertex closest to the center of mass of each parcel (Schaefer-400 7 Network (Schaefer et al., 2018)), b) transcriptional similarity (Pearson’s correlation), defined as the correlation between cortical parcels’ expression profiles across 8,088 stable genes (r > 0.1) derived from the Allen Human Brain Atlas (AHBA) (Hawrylycz et al., 2012), and c) receptor similarity (Pearson’s correlation), defined as the correlation between cortical parcels’ density profiles across 19 receptors/transporters (Dukart et al., 2021; Hansen et al., 2022).

##### Effects of Spatial Autocorrelation on Mapping Performance

Subsequent analyses were conducted to investigate whether mapping accuracy might be driven by spatial autocorrelation.

First, given that spatial autocorrelation might inflate the estimated association between brain maps (Alexander-Bloch et al., 2018; Markello & Misic, 2021), a distance-dependent cross-validation was first performed (Hansen et al., 2021; Hansen et al., 2022). For each cortical region of the Schaefer-400 atlas (source node), the 75% nearest regions to each source node in surface space were selected as the training set (short-range connections), leaving the remaining 25% of regions farthest from the source node for the test set (long-range connections). Mapping accuracy was aggregated across each source node for the training (short-range connections) and test (long-range connections) sets for each subject and compared using *R*^2^, MAE, and RMSE.

A second analysis was conducted to examine whether propagation mapping captures patterns beyond the effect of regional proximity. Permutation tests were performed in three consecutive blocks: cortico-cortical (Hungarian algorithm), subcortical-subcortical (shuffle), and cortico-subcortical (hybrid). First, cortical regions underwent 1,000 permutations using the Hungarian algorithm to generate a null distribution for cortico-cortical propagation edges. Coordinates were defined by selecting the coordinates of the vertex closest to the center of the mass of each parcel, then randomly rotated and reassigned using the Hungarian algorithm, preserving spatial autocorrelation. Second, subcortical-subcortical null distributions were generated by randomly shuffling subcortical regions 1,000 times. Propagation maps were computed for each of these null activity maps to derive null distributions for cortico-cortical and subcortical-subcortical edges. Since cortical and subcortical nulls were generated separately, a hybrid cortical–subcortical null was additionally created by simultaneously rotating cortical regions and shuffling subcortical regions, with propagation edges between cortical and subcortical regions calculated for each permutation. Z-scores for all edges were then computed by comparing observed propagation values to the mean and standard deviation of the combined null distributions, thresholded (p<0.05), and binarized. The difference in mapping accuracy between the observed propagation model and the spatially constrained null models (ΔR^2^) was calculated, and statistical significance was determined using permutation-based one-sample tests.

### 2.3 Study 2

#### 2.3.1 Study sample and Design

A second study was conducted to examine the replicability of predictive accuracy of propagation mapping using resting-state spontaneous brain activity, as well as to validate whether propagation mapping preserves subject’s unique brain organization. To do so, neuroimaging data from 227 healthy individuals (82 females, age range 20-77 years old) were derived from the publicly available Mind-Brain-Body dataset from the Max Planck Institute for Human and Cognitive and Brain Sciences (Babayan et al., 2019). Participants resting-state MRI session for a duration of 15 min 30. Data acquisition and preprocessing can be found elsewhere (Babayan et al., 2019) and in Supplementary Material.

#### 2.3.1 Statistical Analyses

##### Propagation Mapping Performance Using Low-Frequency Oscillations

Propagation mapping was tested using low frequency oscillations of BOLD signal during resting-state. This was captured by amplitude of low frequency fluctuations (ALFF) which assess the strength of spontaneous neural activity within the low frequency band (Zang et al., 2007). ALFF is calculated as the root mean square of BOLD signal at each individual voxel after band-pass filtering (Yang et al., 2007). Although ALFF shows moderate to high test-retest reliability (Zuo et al., 2010), it is also sensitive to physiological noise. Therefore, fractional ALFF (fALFF) was also computed by taking the power of low frequency range divided by the total power of entire frequency range (Zou et al., 2008).

First, the generalizability of propagation mapping at the subject level was evaluated by repeating the Study 1 analyses (i.e., across different parcellations, signal intensities, and spatial distances). To test generalizability of the approach to group-level inferences, one-sample permutation tests was first conducted on ALFF and fALFF regional maps of the 189 subjects who passed quality assurance. These group-level maps were subsequently parcellated. In parallel, a propagation map was generated for each subject, yielding 189 ALFF-based and 189 fALFF-based propagation maps, which were then aggregated at the group level using one-sample permutation tests implemented in the Network-Based Statistics toolbox (Andrew Zalesky, Fornito, & Bullmore, 2010). The rows of the resulting group-level propagation matrices were summed to derive predicted group-level maps, which were compared with the observed group-level (f)ALFF maps using R^2^, MAE, and RMSE.

##### Inter-Individual Variability in Propagation Patterns

Growing evidence suggests that individuals exhibit unique brain activity patterns, often referred to as functional connectome fingerprint (Amico & Goñi, 2018; Finn et al., 2015). Given that the current method relies on normative FC and SC derived from an independent sample (GSP1000), it is possible that subject-specific variability is significantly reduced, which ultimately limit the utility of propagation mapping for studying inter-individual differences. Therefore, analysis, analogous to functional connectome fingerprint (Amico & Goñi, 2018; Finn et al., 2015) were conducted to identify whether propagation mapping indeed reduce subject-specific variance, compared to actual amplitude vectors. Specifically, for each subject (*s*), propagation maps (*PM*) were generated from the two measures of spontaneous brain activity, namely ALFF & fALFF. For each subject, the upper diagonal of the propagation map was stacked into a vector (88,410, edges). A within-subject similarity (*ISelf*) was first defined as the Pearson correlation between subject *s*’s ALFF-based and fALFF-based propagation maps:

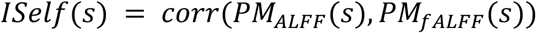

A between-subject similarity (*IOther*) was then defined as the average correlation between subject *s*’s propagation maps (*PM_ALFF_* & *PM_ALFF_*) with those of all other subjects (*i ≠ s*), where *N* is the total number of subjects(S. Stampacchia et al., 2024):

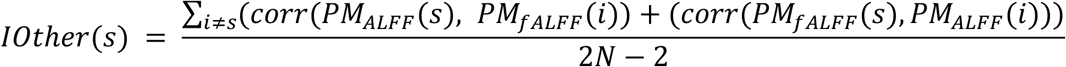

Finally, the brain’s identifiability was measured by two metrics ways, in line with recent work (Amico & Goñi, 2018; Sara Stampacchia et al., 2024). First, brain discriminability (*IDiff*) was quantified as the difference between a subject’s similarity to themselves (*ISelf*) and their similarity to other subjects (*IOthers*). Paired statistical comparisons were then conducted to evaluate differences in subject identifiability between observed amplitude vectors and estimated propagation maps. For each identifiability metric (*ISelf, IOther*, and *IDiff*), within-subject differences were assessed using paired permutation tests (10,000 permutations). Effect sizes were quantified using Cohen’s dz to characterize the magnitude of paired differences.

Next, to assess the contribution of intrinsic connectivity networks to the identifiability, success rate was calculated as the percentage of cases in which the within-subject similarity (*ISelf*, diagonal elements of the matrix) exceeded the between-subject similarity (*IOthers,* off-diagonal elements of the matrix). This metric was computed separately for within- and between-network connections across the 7 Schaefer’s canonical networks and the subcortex.

## 3 Results

### 3.1 Study 1

#### 3.1.1 Propagation Mapping Performance Across Biological Models

Mapping accuracy across different biological models was first evaluated to identify the model that best captures how brain activity propagates throughout the brain. Across six task conditions and five parcellation schemes, the best-fitting model combined the positive edges of FC and SC matrices with unequal weighting, while using the full FC matrix provides the worst fit, as demonstrated by *R*^2^, MAE, and RMSE. More precisely, the best alpha was found by averaging values across task conditions at highest *R*^2^ , and lowest MAE & RMSE for each parcellation scheme. Propagation mapping accuracy was similar across parcellations, with best performance for Schaefer7 (α = 0.7, avg *R*^2^ = 0.95, avg MAE = 0.15; avg RMSE = 0.23), Schaefer17 (α = 0.715, avg *R*^2^ = 0.94, avg MAE = 0.16; avg RMSE = 0.24), and Glasser (α = 0.581, avg *R*^2^ = 0.94, avg MAE = 0.18; avg RMSE = 0.25), and slightly lowest for Gordon (α = 0.61, avg *R*^2^ = 0.93, avg MAE = 0.18; avg RMSE = 0.27), and Desikan-Kiliany (α = 0.554, avg *R*^2^ = 0.93, avg MAE = 0.18; avg RMSE = 0.26) atlases. Mapping accuracy was also highly similar across task conditions (see Figure 2A, and Figures S1-S7).

**Figure 2.**
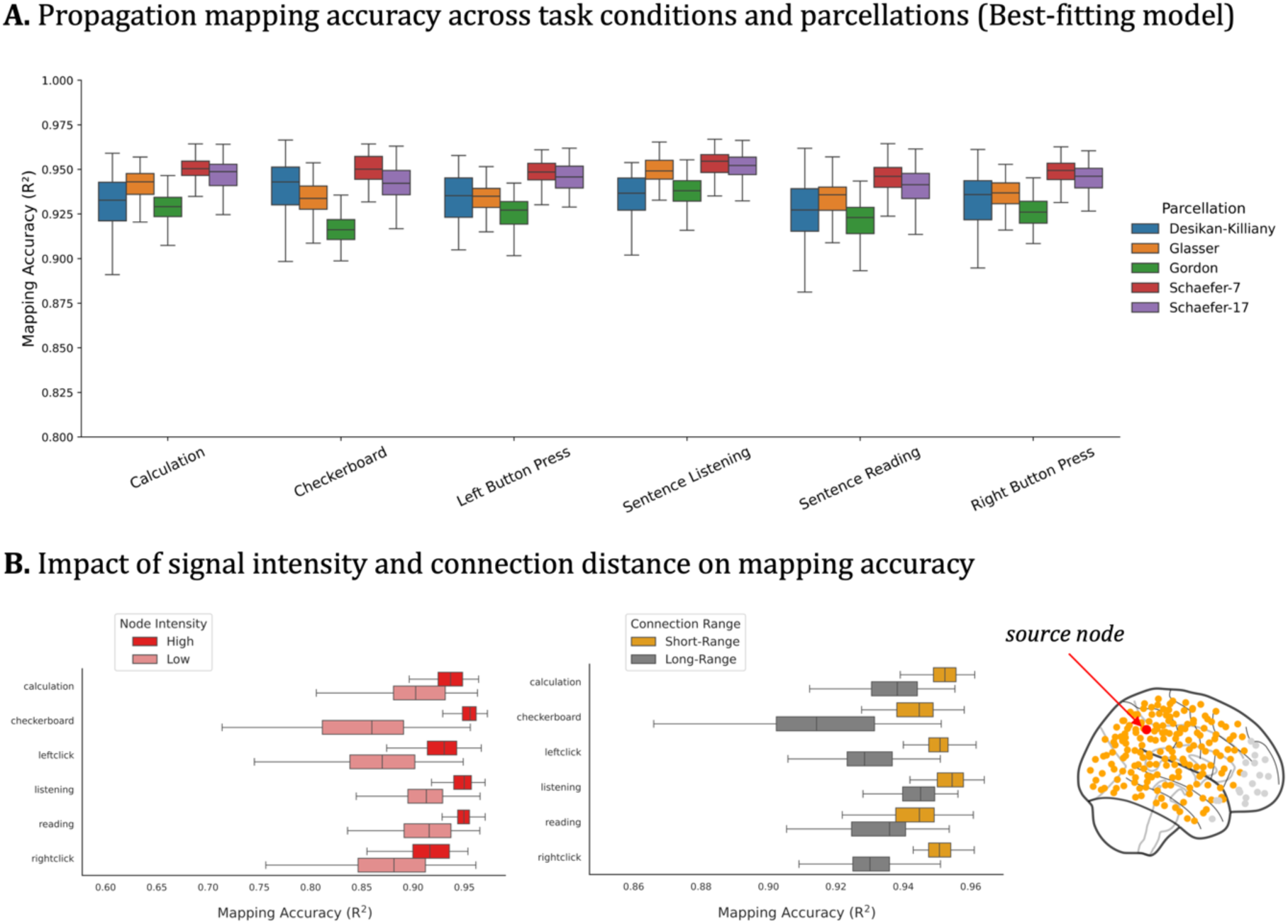
Performance of Propagation Mapping (Study 1). A. Bar graphs show the subject-level mapping accuracy (R2) across parcellation atlases and task contrasts. Across multiple task contrasts, mapping accuracy was the highest using Schaefer’s 7-Network parcellation (avg R2 = 0.95, avg MAE = 0.15, RMSE = 0.23) and the lowest using Gordon parcellation (avg R2 = 0.93, avg MAE = 0.18; avg RMSE = 0.27). B. Subanalyses examined the impact of signal intensity and spatial proximity of brain regions. The left panel shows mapping accuracy when analyses were restricted to nodes in the top 25% highest signal intensity (R^2^ range = 0.92–0.95) and the bottom 25% lowest signal intensity (R^2^ = 0.84–0.91), analyzed separately. C. To test whether performance was driven by short-range connections, distance-based cross-validation was conducted. The training set consisted of the 75% of regions nearest to each source node in surface space, and the test set consisted of the 25% of regions farthest from the source node. Accuracy of propagation mapping was aggregated across each source node iterations for the training (short-range connections, R^2^ range from 0.94 to 0.95) and test (long-range connections, (R^2^ range from 0.91 to 0.94) sets for each subject.

**Figure 3.**
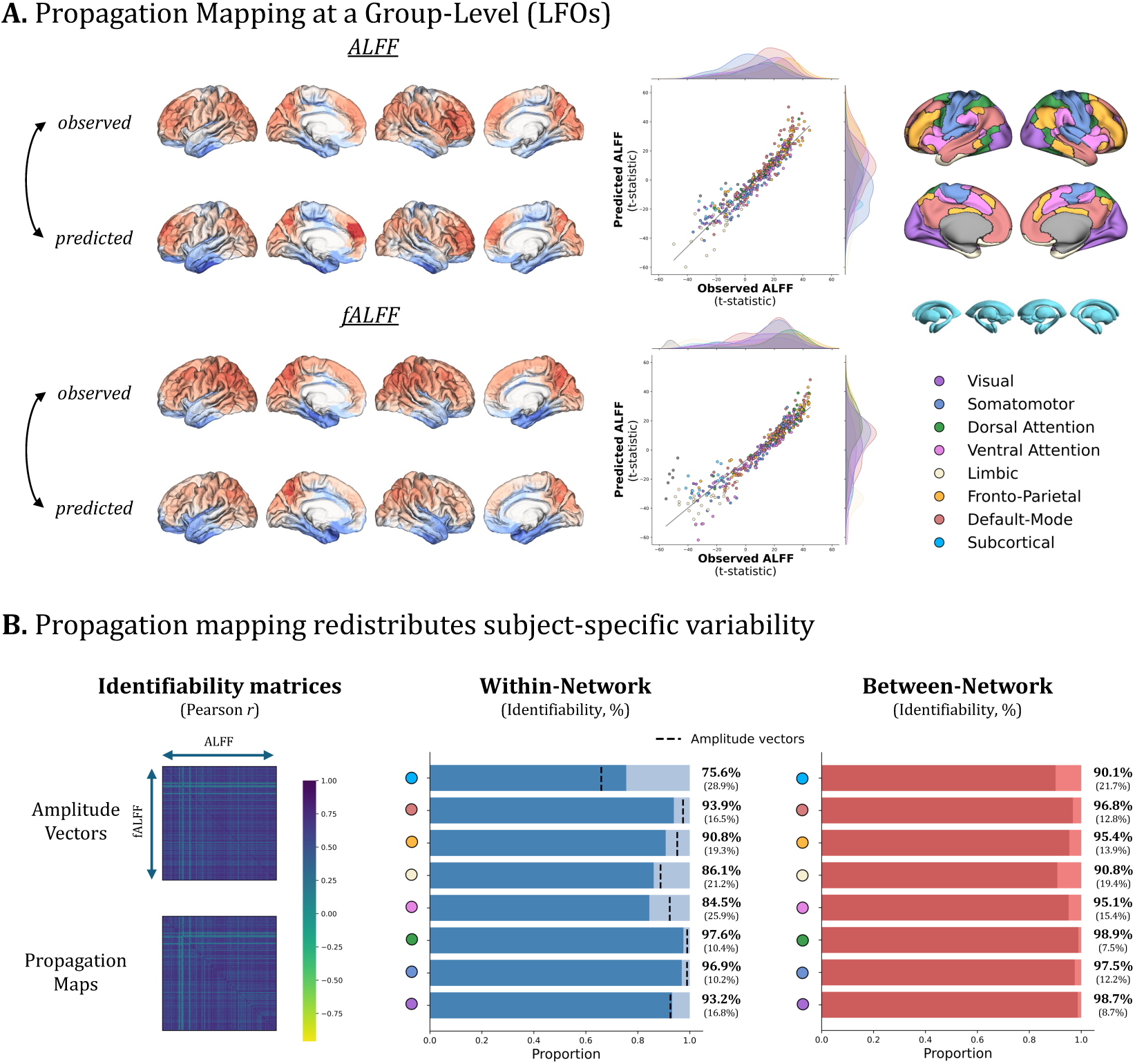
Applicability of Propagation Mapping to resting-state neuroimaging data (Study 2). A. Analyses were conducted to examine the extent to which propagation mapping preserves group-level information. Propagation mapping was first applied to subjects’ resting-state spontaneous brain activity measures, including amplitude of low-frequency fluctuations (ALFF) and its fractional variant (fALFF). Edgewise one-sample t-tests were performed across subjects to generate a group-level propagation map for each measure. Rows of these group-level maps were summed to generate a weighted degree (strength) vector, which was subsequently compared to the original group-level amplitude vector. Results demonstrated that propagation mapping accurately captures group-level information, as reflected by the strong correlation between the predicted and true amplitude vectors. Accuracy for fALFF was reduced in low-amplitude regions, potentially due to normalization by the total power. B. Analyses, analogous to functional connectome fingerprint (Finn et al., 2015) were performed to assess whether propagation maps capture inter-individual variability. Identifiability matrices derived from propagation maps were compared with those from the observed amplitude vectors, allowing direct evaluation of subject-specific information. Results indicate no significant differences in brain discriminality (IDiff) between the two approaches (Cohen’s dz = 0.10, p = 0.169), suggesting that propagation mapping redistributes inter-individual variability along propagation routes. LFOs = Low Frequency Oscillations.

To ensure that mapping accuracy was not driven by differences in signal intensity, analyses were repeated separately for regions with the highest (top 25%) and lowest (bottom 25%) signal intensity. Across task conditions, propagation mapping achieved higher accuracy when restricted to high-intensity regions (R^2^ = 0.92–0.95), but remained relatively accurate even in low-intensity regions (R^2^ = 0.84–0.91)(Figure 2B & Figures S8). Moreover, the mapping accuracy of the best biological model for each of the parcellation was highly similar to when propagation mapping was conducted on 1,000 randomly selected fMRI brain maps: Schaefer7 (Δ*R*^2^=0.001; ΔMAE=0.006; ΔRMSE=0.006), Schaefer17 (Δ*R*^2^=0.005; ΔMAE=0.008; ΔRMSE=0.013), Glasser (Δ*R*^2^=0.006; ΔMAE=0.012; ΔRMSE=0.014); Gordon (Δ*R*^2^=0.002; ΔMAE=0.007; ΔRMSE=0.007); Desikan (Δ*R*^2^=0.011; ΔMAE=0.012; ΔRMSE=0.024) (Figure S9).

Post hoc analyses were further conducted to improve interpretability of the role of SC in identifying propagation routes relative to FC. Spatial distance between brain regions, transcriptomics similarity, and receptor similarity were computed. Together, these variables explained 21.0% of the variance in the FC matrix and 18.9% of the variance in the SC matrix. Dominance analyses further revealed differential relative contributions (Figure S10): in FC, spatial proximity accounted for 53.9% of the explained variance, whereas in SC it accounted for 40.0%. In contrast, receptor similarity contributed a larger proportion of variance in SC (50.2%) than in FC (37.2%), while transcriptomic similarity contributed similarly in both models (approximately 9–10%). These findings suggest that SC may indeed help constrain FC to better capture underlying task-evoked propagation routes.

#### 3.1.2 Effects of Spatial Autocorrelation on Mapping Performance

To ensure that the mapping accuracy was not solely driven by spatial autocorrelation, two analyses were carried out. First, distance-dependent cross-validation was conducted by identifying the 75% nearest regions (short-range connections) and the 25% farthest regions (long-range connections) for each source node (parcels) and aggregated. Results showed that while short range connections have the highest mapping accuracy across different task conditions (*R*^2^ range from 0.94 to 0.95; MAE: 0.14-0.16; RMSE: 0.21-0.23), long-range connections also contribute to the propagation mapping (*R*^2^: 0.91-0.94; MAE: 0.16-0.17; RMSE: 0.22-0.24) (Figure 2B, Right Panel, and Figure S11).

A spatially constrained null was applied to confirm that propagation effects reflect network topology beyond what is explained by spatial proximity. Across all task conditions, propagation maps showed significantly higher mapping accuracy than the null (ΔR^2^ = 0.92–0.94, p < 0.001; ΔMAE = 0.56–0.62; ΔRMSE = 0.75–0.77). Importantly, when considering only edges that survived statistical significance after the spatial null (p < 0.05), propagation mapping still explained a substantial portion of variance in brain activity (R^2^ = 0.58–0.66), despite some error (MAE = 0.46–0.53; RMSE = 0.58–0.65).

### 3.2 Study 2

#### 3.2.1 Propagation Mapping Performance Using Low-Frequency Oscillations

An independent sample (n = 189) was used to test the replicability of propagation mapping’s performance using low-frequency oscillations of BOLD signal during restint-state. Propagation mapping showed slightly lower accuracy in fALFF (*R*^2^ range 0.90-0.92) relative to ALFF (*R*^2^ range 0.93-0.95) across parcellations (Figure S12). Similar performance was found between short- and long-range connections for both ALFF (*R*^2^_SHORT_=0.94; *R*^2^_LONG_=0.93) and fALFF (*R*^2^_SHORT_=0.93; *R*^2^_LONG_=0.92) (Figure S13). However, regions with below-average amplitude (z < 0) contributed less to the mapping accuracy, especially for fALFF (*R*^2^_HIGH_=0.90; *R*^2^_LOW_=0.35), relative to ALFF (*R*^2^_HIGH_=0.88; *R*^2^_LOW_=0.73) (Figure S14). These indicates that low-amplitude regions in fALFF may carry little signal variance due to its normalization by total power, whereas ALFF preserves absolute amplitude, allowing propagation mapping to perform well even in weaker regions. Indeed, the effect disappeared when using absolute amplitude scaling for defining high-low intensity regions in both ALFF (*R*^2^_HIGH_=0.95; *R*^2^_LOW_=0.85) and fALFF (*R*^2^_HIGH_=0.90; *R*^2^_LOW_=0.85) (Figure S15).

At the group level, the degree strength of propagation maps showed slightly lower accuracy in mapping observed group-level amplitude t-maps compared to the subject-level performance reported in Study 1 (ALFF: R^2^ = 0.89, MAE = 4.46, and RMSE = 6.09; fALFF: R^2^ = 0.88, MAE = 5.37, and RMSE = 8.30). These results indicate that propagation mapping provides relatively stable mapping performance across different levels of analysis.

#### Inter-Individual Variability in Propagation Patterns

Given that the current model relies on normative connectomes estimated from a large sample of healthy subjects, it is possible that this approach smooths out subject-specific variance, potentially limiting the usefulness of propagation mapping for studying inter-individual differences. Therefore, analyses analogous to functional connectome fingerprinting were conducted to examine differences in identifiability between subjects’ amplitude vectors and their propagation maps. To do so, each subject’s ALFF- and fALFF-based propagation maps were correlated with themselves (ISelf) and with other subjects (IOthers) to create an identifiability matrix (Figure 2B). The same procedure was applied to subjects’ ALFF and fALFF amplitude vectors to enable comparability. Paired t-tests revealed that individuals showed greater self-similarity (ISelf) in amplitude vectors than in propagation maps (Amplitude mean = 0.77 vs. Propagation mean = 0.75; Cohen’s dz = 0.53, p < 0.001). Conversely, propagation maps exhibited lower between-subject similarity (IOther) relative to amplitude vectors (Amplitude mean = 0.59 vs. Propagation mean = 0.58; Cohen’s dz = 0.18, p = 0.015). Overall, the brain discriminability (IDiff), as measured by the difference between self- and other-similarity, did not significantly differ between the two approaches (Amplitude mean = 0.181 vs. Propagation mean = 0.176; Cohen’s dz = 0.10, p = 0.169), suggesting that propagation mapping does not suppress inter-individual variability but rather redistributes it along propagation routes.

Despite slightly lower identification success rates for propagation maps compared to amplitude vectors (Amplitude mean = 98.5% vs. Propagation mean = 94.6%; Cohen’s d = 0.39, p < 0.001), propagation mapping still achieves identification accuracy comparable to those reported in the seminal study on functional connectome fingerprinting (Finn et al., 2015). As shown in Figure 2B, identification accuracy (success rate) was generally higher for between-network connectivity compared to within-network connectivity, as also reported in functional connectome fingerprinting studies (St-Onge et al., 2023). Indeed, near perfect identification accuracy was found for between-network propagation from the Dorsal Attention (98.9%, SD=7.5%), Visual (98.7%, SD=8.7%), and the Somatomotor (97.5%, SD=12.2%), and networks.

## 5 Discussion

The goal of the current study was to introduce and validate *propagation mapping*, a method designed to model the propagation of task-evoked brain activity throughout the whole-brain. Findings of the current study underscore that regional activity and connectivity are not mutually exclusive measures but fundamentally interdependent, as previously demonstrated by extensive work on activity flow mapping (Cocuzza et al., 2024; Cole et al., 2016; Cole et al., 2021; Sanchez-Romero et al., 2023). Importantly, while resting-state FC can indeed predict task-evoked activity (Cole et al., 2016), the mapping accuracy of task-evoked activity substantially increased when SC was included as a neuroanatomical constraint to establish the propagation routes. The performance of the current method was consistent across multiple task contrasts and parcellation atlases, and was robust to differences in signal intensity and regional spatial proximity. Furthermore, despite relying on normative connectomes, propagation mapping does not significantly reduce individual differences, but rather redistributes subject-specific variability along whole-brain topological routes. Overall, propagation mapping is a reliable and easy to use tool to capture how task-evoked activity propagates throughout the brain, opening new possibilities for neuroimaging research.

A significant novelty of the current approach is the ability to capture how regional variations in brain activity propagate throughout the whole brain. The method achieved high mapping accuracy, suggesting that the model successfully capture how task-evoked activity propagates. This can be attributed to two key factors: (1) the stability and robustness of connectivity estimates obtained from a large cohort of healthy individuals, and (2) the incorporation of structural covariance as an anatomical constraint. A series of biological models were first tested to assess the performance of propagation mapping. Findings suggested that mapping accuracy was highest when positive edges of SC and FC matrices were combined, with a larger weighting assigned to SC relative to FC. First, as methods for reconstructing brain connectomes are prone to erroneously infer connectivity between disconnected pairs of regions (van den Heuvel et al., 2017; Zalesky et al., 2016), controlling for spurious connections is crucial. For instance, previous research has shown that SC correlates with resting-state FC (Alexander-Bloch et al., 2013; Mechelli et al., 2005), suggesting that SC may reveal biologically meaningful patterns. In line with this, the post hoc dominance analyses conducted in the current study supported the interpretation that SC serves as a biological constraint on FC to identify reliable propagation routes. Nevertheless, future work may help clarify the extent to which these propagation routes reflect underlying white matter anatomical connections. Second, it is worth noting that positive edges alone contributed substantially to mapping accuracy compared to the full connectivity range, indicating that negative edges (or anti-correlations) play little role in explaining the propagation of task-evoked activity. This aligns with prior work showing greater reliability of positive compared to negative correlations (Shehzad et al., 2009) and supports the idea that positive connections form the core architecture underlying brain spatial organization (Cole et al., 2014; Gratton et al., 2018; Smith et al., 2009).

Despite that the current method appears to effectively capture how task-evoked activity propagates throughout the brain, several factors may moderate the findings. First, propagation mapping performance may be affected in regions with low signal variance, where MRI artifacts or drop-out could artificially reduce variability and influence connectivity estimates. Therefore, sub-analyses were conducted to evaluate the stability of propagation mapping by separately restricting the analysis to nodes with the top 25% highest and bottom 25% lowest signal intensity. In task-evoked activity, the method performed slightly better when restricting to high-intensity regions (R^2^ ranging from 0.92 to 0.95), relative to low-intensity regions (R^2^ ranging from 0.84 to 0.91). This pattern was similarly observed in resting-state ALFF, suggesting that the method is generally robust to low-variance regions. However, mapping accuracy in low-intensity regions dropped substantially when examining resting-state fALFF (R^2^ = 0.35). One possible explanation is that propagation mapping requires sufficient signal amplitude for the model to detect propagation across the weighted connectome. Normalization in fALFF may reduce signal variance in low-amplitude regions compared with ALFF, which preserves absolute amplitude. Second, spatial proximity between regions is known to inflate neuroimaging findings when spatial autocorrelation is not accounted for (Alexander-Bloch et al., 2018; Burt, Helmer, Shinn, Anticevic, & Murray, 2020; Markello & Misic, 2021). Through a distance-dependent cross-validation, findings of the current study provide the evidence that restricting the analyses to the 25% farthest regions of the source nodes revealed largely similar findings (*R*^2^: 0.91-0.94 in long-range connections, versus *R*^2^ range from 0.94 to 0.95 in short-range connections). To further explore the influence of spatial proximity on mapping performance, spatial autocorrelation–preserving nulls were generated. Results indicate that spatial autocorrelation alone does not fully account for the observed propagation patterns (ΔR^2^ = 0.92–0.94, p < 0.001). Even when analyses were restricted to region-to-region connections that exceeded the effect of regional proximity (p < 0.05), propagation mapping continued to explain a substantial portion of variance in task-evoked activity (R^2^ = 0.58–0.66). As shown by the post hoc dominance analyses, it is possible that the addition of SC significantly reduced the impact of spatial proximity. Overall, propagation mapping offers powerful and easy-to-use tool for mapping propagation patterns which expands the potential avenues for neuroimaging research. For example, the current method does not require resting- state data to identify propagation routes, as it leverages normative connectomes. Importantly, it demonstrates good discriminability between subjects, highlighting the potential of propagation mapping for studying inter-individual variability. Notably, compared with other methods that rely on normative connectomes (Fox, 2018), which have been recently criticized for their limited variability between brain maps (van den Heuvel et al., 2026), propagation mapping may provide a valuable alternative. Similarly, it can be used for graph metrics and connectome-based predictive modeling (Shen et al., 2017). Other potential benefits of propagation mapping include generating novel research questions from static images without requiring time-series data. Indeed, although some group-level approaches based on inter-individual covariance have been proposed (Alexander-Bloch et al., 2013; Huang et al., 2010), connectivity-based inferences at the subject level remain challenging for many modalities (e.g., positron emission tomography, structural imaging, imaging transcriptomics). Propagation mapping also provides residual measures, enabling the identification of regions that deviate from normative expectations and offering novel insights into atypical brain activity. Additionally, computational lesioning could be applied to quantify the relative importance of individual nodes in the propagation of task-evoked activity. In sum, it is future applications of this method may further advance research in neurological and psychiatric disorders.

The method described in the current study provides a direct and easy-to-use alternative to characterize the propagation of brain activity. This method attempts to reconcile localizationist and connectionist perspectives on the conceptualization of the brain, with the aim of generating new discoveries in neuroscience and psychiatric imaging. Despite the strengths of this method, several limitations should be acknowledged. First, propagation mapping does not achieve perfect accuracy. Therefore, researchers should consider accounting for variability in mapping accuracy when studying inter-individual differences. Second, the current method relies on normative data estimated from the GSP1000 dataset, which provides a fast and easy way to capture propagation patterns. While the method was originally developed as an alternative to overcome the noisy signals and short acquisition times of individual resting-state scans, subject-level data nonetheless provide a more idiosyncratic characterization of both adaptive (e.g., compensation) and maladaptive (e.g., diaschisis) propagation patterns. Future studies could explore the benefits of integrating prior knowledge of covariance distributions (Rahim, Thirion, & Varoquaux, 2019) with individual resting-state data to improve subject-specific propagation maps. Third, the method relies on a bivariate correlation approach, meaning that while it captures the total shared variance between regions, the directionality of signal propagation cannot be determined. This approach is consistent with anatomical evidence of bidirectional projections among many brain regions (Haber, 2011; Markov & Kennedy, 2013), and provides an interpretable measure of network-wide propagation without relying on the stringent assumptions required by models designed to isolate unique regional influences (Friston, 2011; Sanchez-Romero et al., 2023). Future work applying alternative models to estimate FC and SC (e.g., multiple regression (Cole et al., 2016), and/or multivariate distance correlation (Geerligs, Cam, & Henson, 2016) may offer further insight into the causal directionality of propagation flow.

## 6 Conclusion

The current study introduced propagation mapping as an extension of activity flow mapping, designed to model how brain activity propagates throughout the whole brain. By integrating normative connectomes, the method achieved robust mapping performance across both task-evoked and spontaneous brain activity. Performance was stable across task conditions, parcellation atlases, levels of analysis, signal intensity, and spatial proximity of brain regions. Importantly, propagation mapping demonstrated strong discriminability between subjects, underscoring its potential for studying inter-individual variability. Overall, this framework provides a valuable tool for advancing scientific discovery in neurological and psychiatric disorders.

## Acknowledgement

I would like to thank the reviewers for their valuable insights and constructive feedback, which have strengthened both the methodology and the manuscript. I also want to thank Dr. Stéphane De Brito for his incommensurable support over the past years, which made this possible.

## Author Contribution

J.R.D. designed the study, performed all analyses, and wrote and approved the final version of the manuscript.

## Data Availability Statement

The Brain Genomics Superstruct Project (https://dataverse.harvard.edu/dataverse/GSP), Mind-Brain-Body dataset from the Max Planck Institute for Human and Cognitive and Brain Sciences (https://fcon_1000.projects.nitrc.org/indi/retro/MPI_LEMON.html), and Brainomics/Localizer dataset (https://osf.io/vhtf6/) are openly available.

## Code Availability Statement

Propagation Mapping Toolbox is a user-friendly toolbox that was made freely available on Streamlit Cloud (https://propagation-mapping.streamlit.app/) or locally (https://github.com/JulDugre/Propagation-Mapping).

