## Supplementary Material for "Propagation Mapping: A Framework for Modeling Whole-Brain Propagation Patterns of Task-Evoked Activity"

### Propagation Mapping:


Jules R. Dugré, PhD [1];

[1] Department of Psychology, College of Literature, Science, and the Arts, University of Michigan, USA.

**Corresponding authors**
Jules Roger Dugré, PhD
Department of Psychology,
College of Literature, Science, and the Arts,
University of Michigan, Ann Arbor, MI, USA 48109


### Table of Contents


***Supplementary Methods. Neuroimaging Data Acquisition*** ... ***4***

**Genomic Superstruct Project (GSP1000) Dataset** ... **4**

**Brainomics/Localizer Dataset** ... **4**

**Mind-Brain-Body Dataset** ... **5**

***Supplementary Methods. Neuroimaging Preprocessing*** ... ***6***

***Supplementary Figures*** ... ***7***

**Figure S1. Determining the Optimal α from Mapping Accuracy ($R^2$) Across Task Conditions and Parcellations (Study 1).** ... **7**

**Figure S2. Determining the Optimal α from Mapping Errors (MAE) Across Task Conditions and Parcellations (Study 1).** ... **8**

**Figure S3. Determining the Optimal α from Mapping Errors (RMSE) Across Task Conditions and Parcellations (Study 1).** ... **9**

**Figure S4. Comparison of Mapping Accuracy ($R^2$) Across Multiple Biological Models, Tasks, and Parcellations (Study 1)** ... **10**

**Figure S5. Comparison of Mapping Errors (MAE) Across Multiple Biological Models, Tasks, and Parcellations (Study 1)** ... **11**

**Figure S6. Comparison of Mapping Errors (RMSE) Across Multiple Biological Models, Tasks, and Parcellations (Study 1)** ... **12**

**Figure S7. Mapping Errors (MAE, RMSE) of the Best Fitting Model for Each Parcellations by Task Conditions (Study 1).** ... **13**

**Figure S8. Impact of signal intensity on Mapping Errors (MAE, RMSE, Study 1)** ... **14**

**Figure S9. Mapping Accuracy ($R^2$) & Errors (MAE, RMSE) of the best fitting propagation model on 1,000 randomly selected functional MRI brain images.** ... **15**

**Figure S10. Post hoc Dominance Analyses on Contributors to Structural Covariance and Functional Connectivity.** ... **16**

**Figure S11. Impact of spatial proximity on Mapping Errors (Study 1).** ... **17**

**Figure S12. Mapping accuracy and errors in an independent sample using the amplitude of low-frequency oscillations during resting-state (Study 2)** ... **18**

**Figure S13. Impact of spatial proximity on mapping accuracy and errors (Study 2).** ... **19**

**Figure S14. Impact of Signal Intensity on Mapping Accuracy and Errors (MAE, RMSE, Study 2).** ... **20**

**Figure S15. Impact of Signal Intensity on Mapping Accuracy and Errors (MAE, RMSE, Study 2)**.......21

***References*** .......22

#### **Supplementary Methods.** Neuroimaging Data Acquisition

##### Genomic Superstruct Project (GSP1000) Dataset

GSP1000 cohort contains a fully 1:1 M:F matched dataset of one thousand human participants ages 18 to 35 years that were scanned with a Siemens Tim Trio 3T scanners (Siemens Healthcare, Erlangen, Germany) at Harvard University and Massachusetts General Hospital using the vendor-supplied 12-channel phased-array head coil. Participants were matched based on age and sex. High-resolution (1.2 mm isotropic) multi-echo MPRAGE sequences were acquired for a duration of 2 min 12s, with sagittal acquisition orientation, 144 slices, TR=2200 ms, TE=1.5/3.4/5.2/7.0 ms, FA= 7°, TI=1100 ms, Resolution =1.2×1.2×1.2 mm. For functional data, subjects were instructed to remain still, stay awake, and keep their eyes open. EPI parameters were as follows: repetition time (TR) = 3,000 ms, echo time (TE) = 30 ms, flip angle (FA) = 85°, 3 × 3 × 3-mm voxels, field of view (FOV) = 216, and 47 axial slices collected with interleaved acquisition and no gap between slices. Each functional run lasted 6.2 min (124 time points). One or two runs were acquired per subject (average of 1.7 runs).

##### Brainomics/Localizer Dataset

Functional images were acquired on a 3T Brucker scanner using an EPI sequence (TR = 2400 ms, TE = 30 ms, matrix size = 64 × 64, FOV = 24 cm × 24 cm). Each volume consisted of 34 slices of 4 mm thickness. Anatomical T1 images were acquired with a spatial resolution of 1 × 1 × 1.2 mm. Data were pre-processed using SPM2 software in Matlab environment according to the following procedure: slice timing, subject motion estimation and correction by realignment, coregistration of the anatomical image to the MNI template, spatial normalization of functional images (resampled voxel size = 3 × 3 × 3 mm) and smoothing (5 mm FWHM). Each voxel time series was fitted with a linear combination of the canonical hemodynamic response function and its temporal derivative. A temporal high pass filter was applied (cutoff 128 sec. and AR(1) whitening).

#### Mind-Brain-Body Dataset

Two hundred and twenty-seven human participants were scanned with a Siemens Magnetom Verio 3 Tesla scanner (Siemens Healthcare GmbH, Erlangen, Germany) equipped with a 32-channel head coil. MP2RAGE sequences were acquired for a duration of 8 min 22 s, with sagittal acquisition orientation, one 3D volume with 176 slices, TR=5000 ms, TE=2.92 ms, TI1=700 ms, TI2=2500 ms, FA1=4°, FA2=5°, pre-scan normalization, echo spacing=6.9 ms, bandwidth=240 Hz/pixel, FOV=256 mm, voxel size= 1 mm isotropic, GRAPPA acceleration factor 3. Four rs-fMRI scans were acquired in axial orientation using T2∗-weighted gradient-echo echo planar imaging (GE-EPI) with multiband acceleration, sensitive to blood oxygen level-dependent (BOLD) contrast. Sequences were identical across the four runs, they differ in alternating slice orientation and phase-encoding direction, to vary the spatial distribution of distortions and signal loss. The y-axis was aligned parallel to the AC-PC axis for runs 1 and 2, and parallel to orbitofrontal cortex for runs 2 and 4. The phase-encoding direction was A–P for runs 1 and 3, and P–A for runs 2 and 4. Further parameters were set as follows for all four runs: voxel size=2.3 mm isotropic, FOV=202×202 mm2, imaging matrix=88×88, 64 slices with 2.3 mm thickness, TR=1400 ms, TE=39.4 ms, flip angle=69°, echo spacing=0.67 ms, bandwidth=1776 Hz/Px, partial fourier 7/8, no pre-scan normalization, multiband acceleration factor=4, 657 volumes, duration=15 min 30 s. During the resting-state scans, participants were instructed to remain awake with their eyes open and to fixate on a crosshair.

#### **Supplementary Methods.** Neuroimaging Preprocessing

Functional images of the Mind-Brain-Body Dataset were realigned, corrected for motion artifacts [1] with the Artifact Detection Tool, setting a threshold of 0.9 mm subject ART's composite motion and a global signal threshold of Z = 5) with the implemented in CONN Toolbox [2], bandpass filtered (0.008 Hz < f < 0.09 Hz) and co-registered to the corresponding anatomical image. The anatomical images were segmented (into grey matter, white matter, and cerebrospinal fluid) and normalized to the Montreal Neurological Institute (MNI) stereotaxic space. Functional images were then normalized based on structural data, spatially smoothed with a 6 mm full-width-at-half-maximum (FWHM) 3D isotropic Gaussian kernel and resampled to 2 $mm^3$ voxels. For the preprocessing, the anatomical component-based noise correction method (aCompCor strategy, [3]), was employed to remove confounding effects from the BOLD time series, such as the physiological noise originating from the white matter and cerebrospinal fluid. This method was found to increase the validity and sensitivity of analyses [4]. Subjects with less than 75% of the original data points remaining after scrubbing, or with high mean motion after scrubbing (>0.25 mm framewise displacement), were excluded. After these exclusions, a final set of 189 subjects was included in the analyses.

### Supplementary Figures

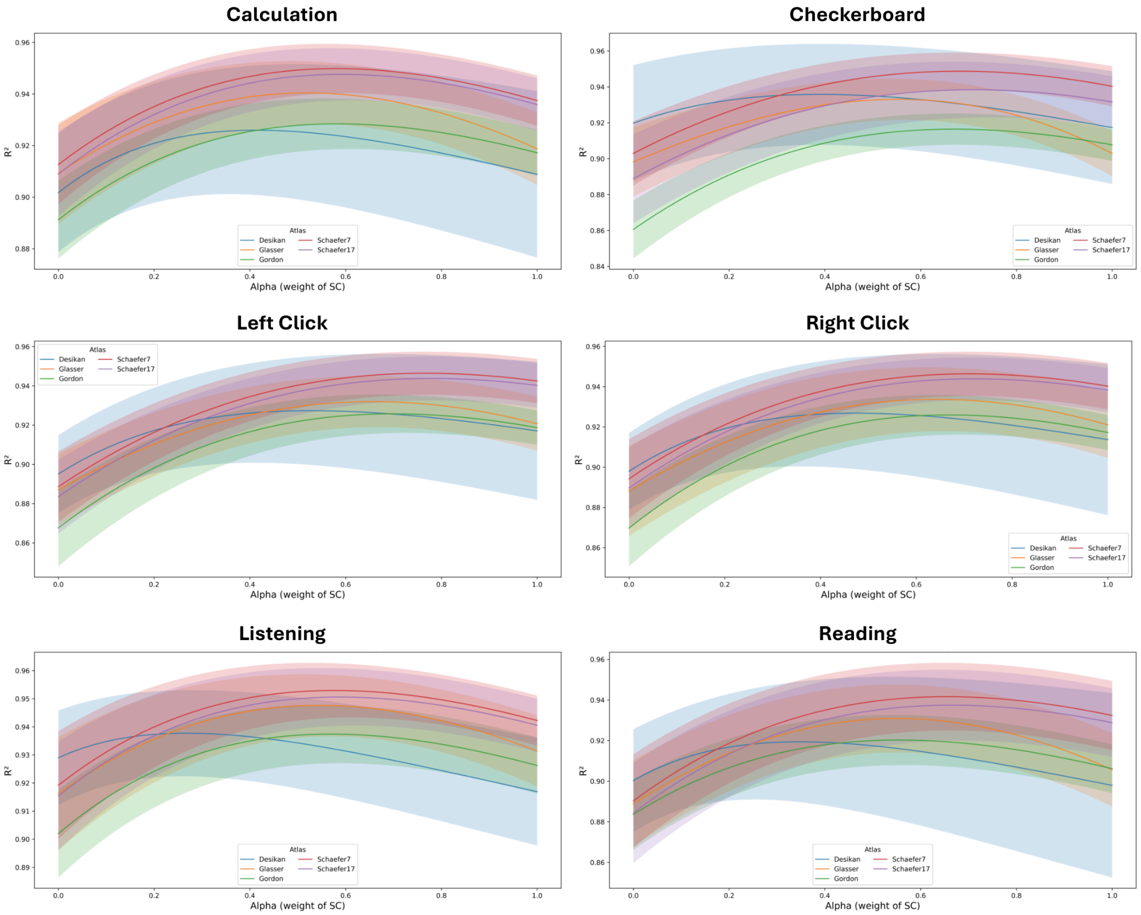


**Figure S1.** Determining the Optimal α from Mapping Accuracy ($R^2$) Across Task Conditions and Parcellations (Study 1).

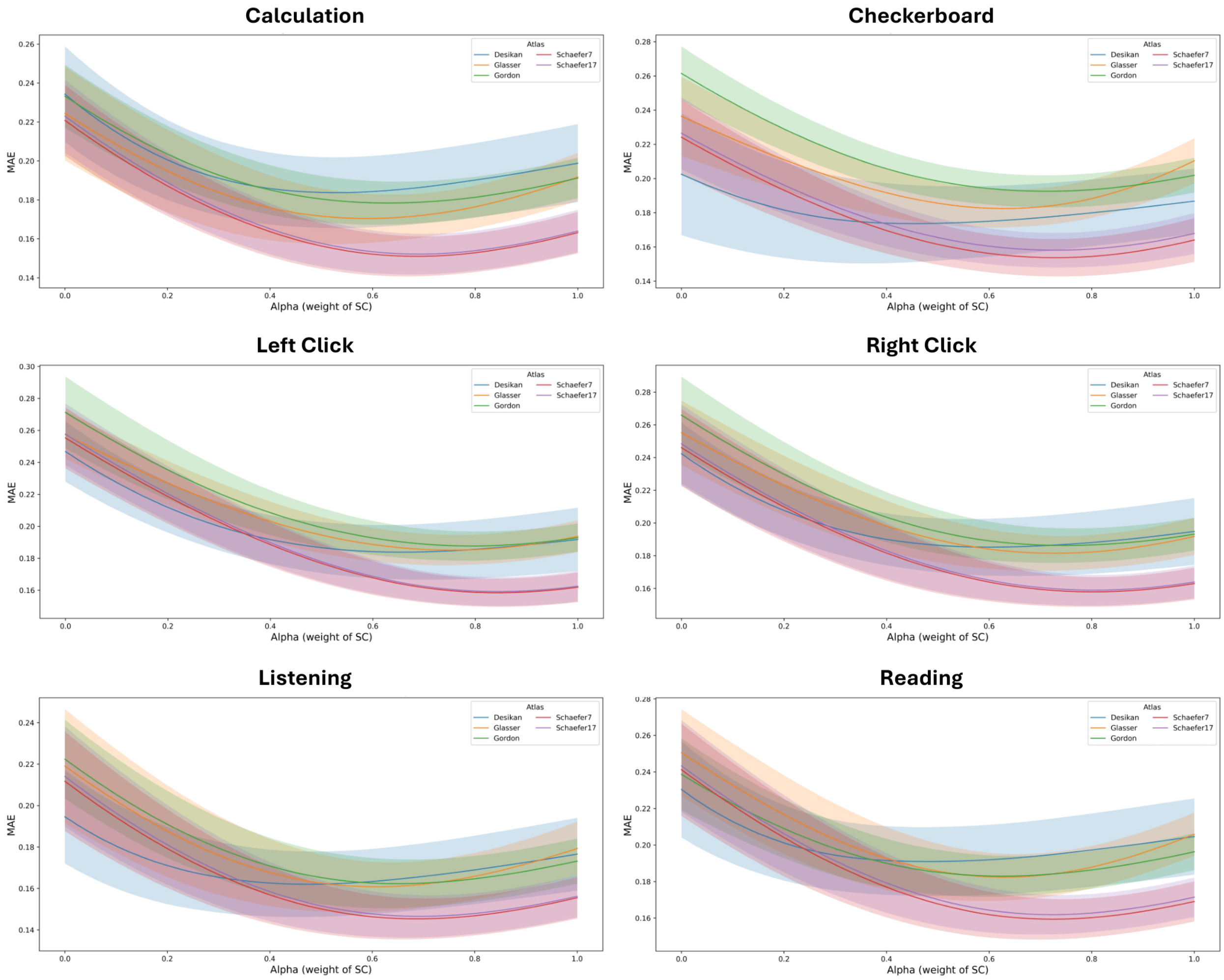


**Figure S2.** Determining the Optimal α from Mapping Errors (MAE) Across Task Conditions and Parcellations (Study 1).

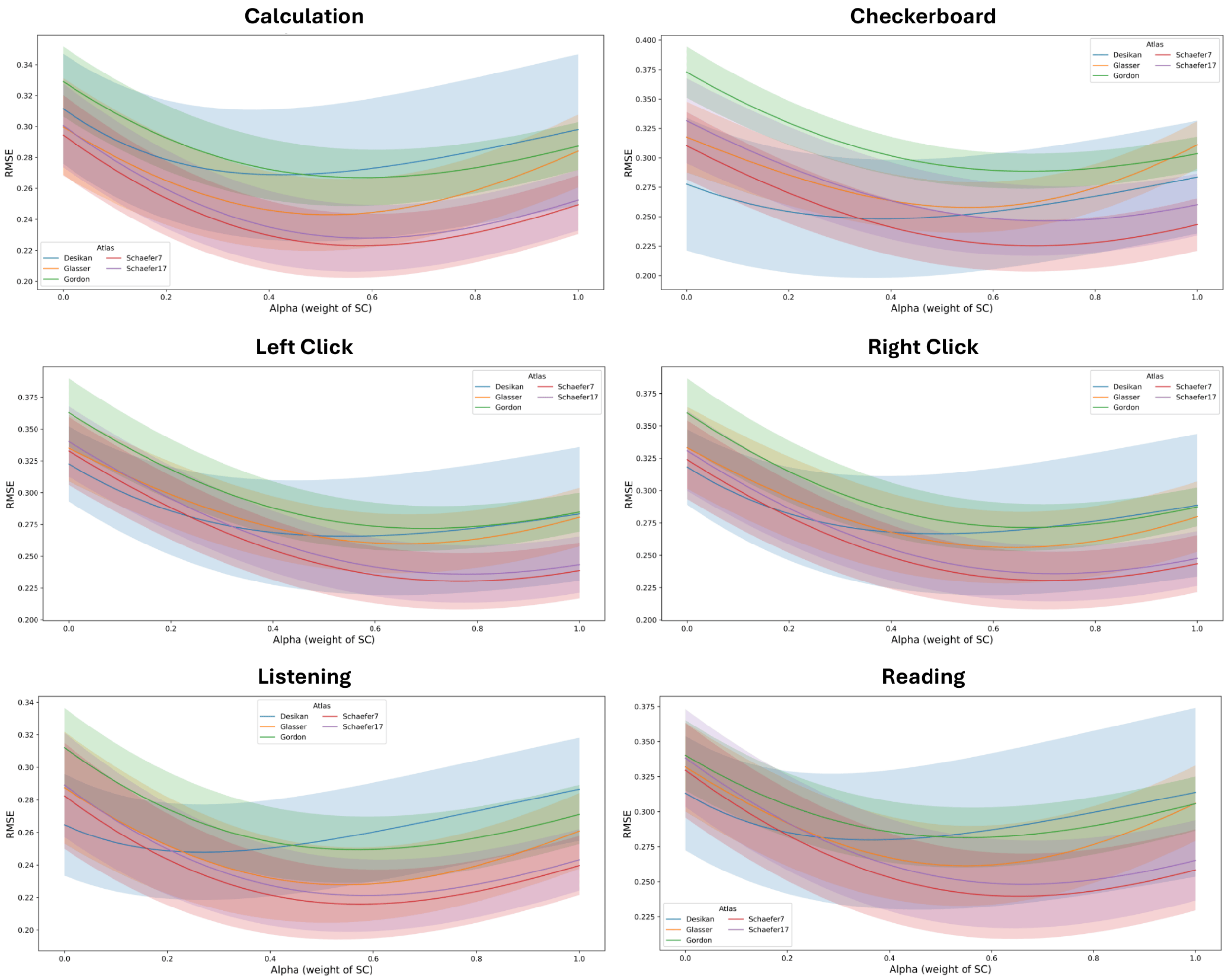


**Figure S3.** Determining the Optimal α from Mapping Errors (RMSE) Across Task Conditions and Parcellations (Study 1).

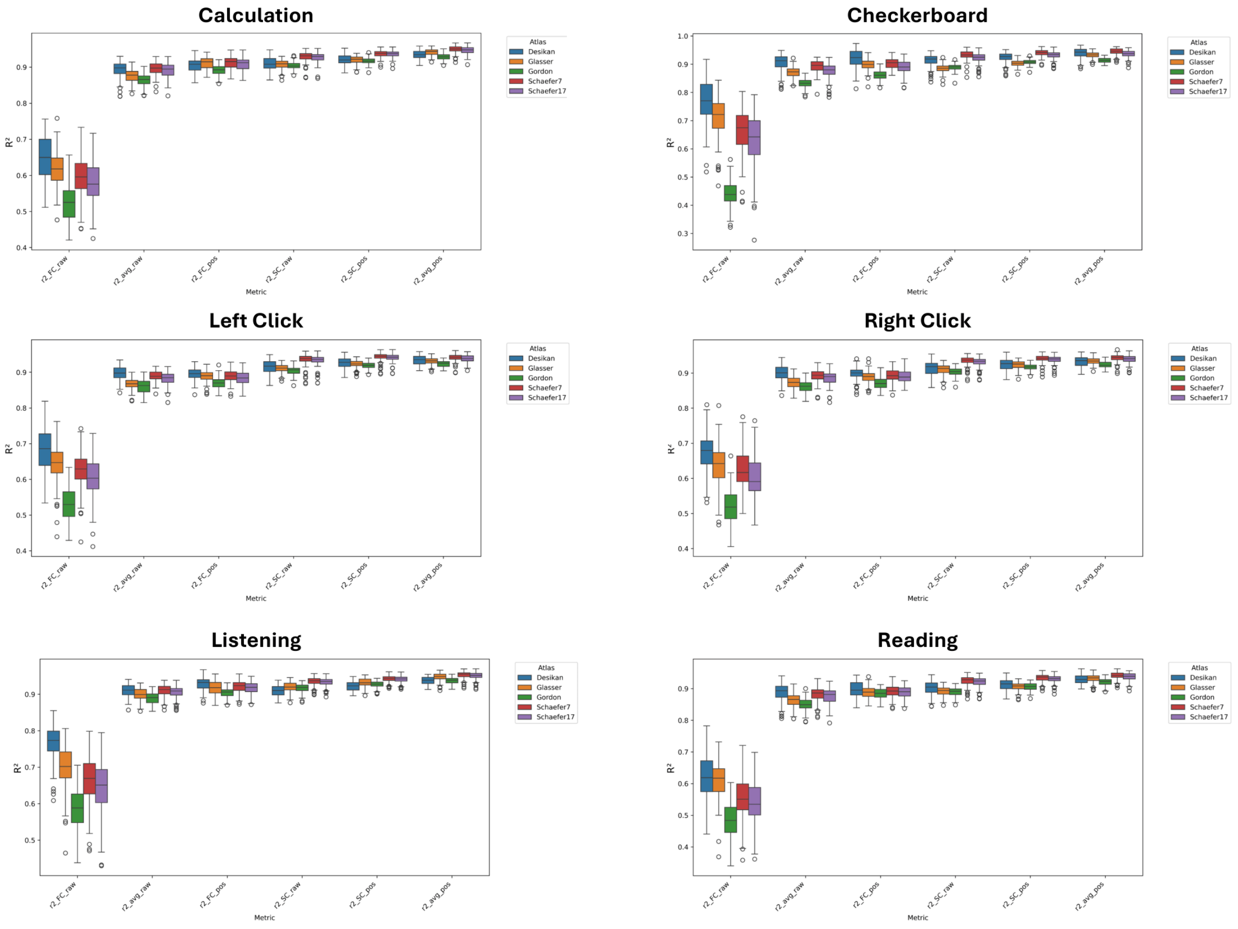


**Figure S4.** Comparison of Mapping Accuracy ($R^2$) Across Multiple Biological Models, Tasks, and Parcellations (Study 1)

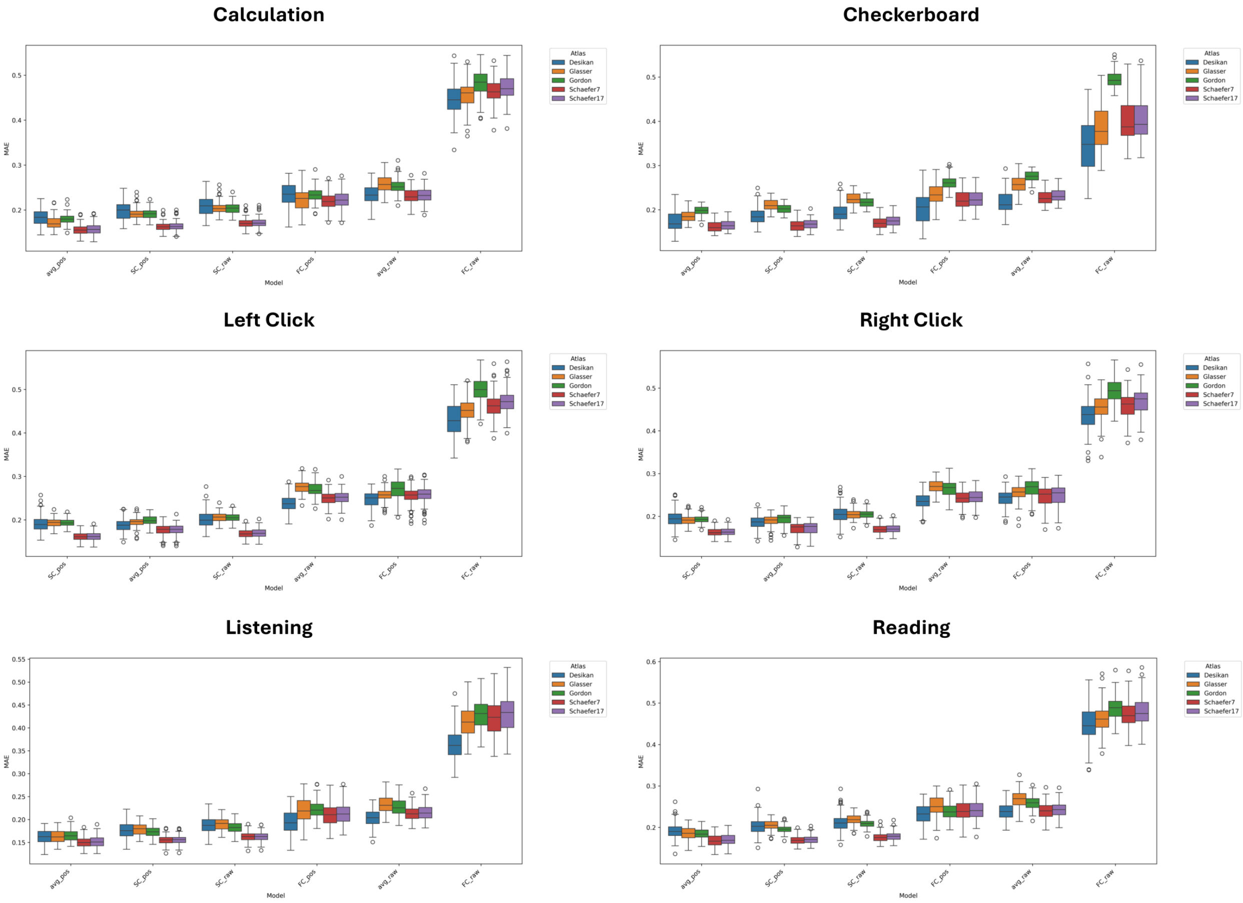


**Figure S5.** Comparison of Mapping Errors (MAE) Across Multiple Biological Models, Tasks, and Parcellations (Study 1)

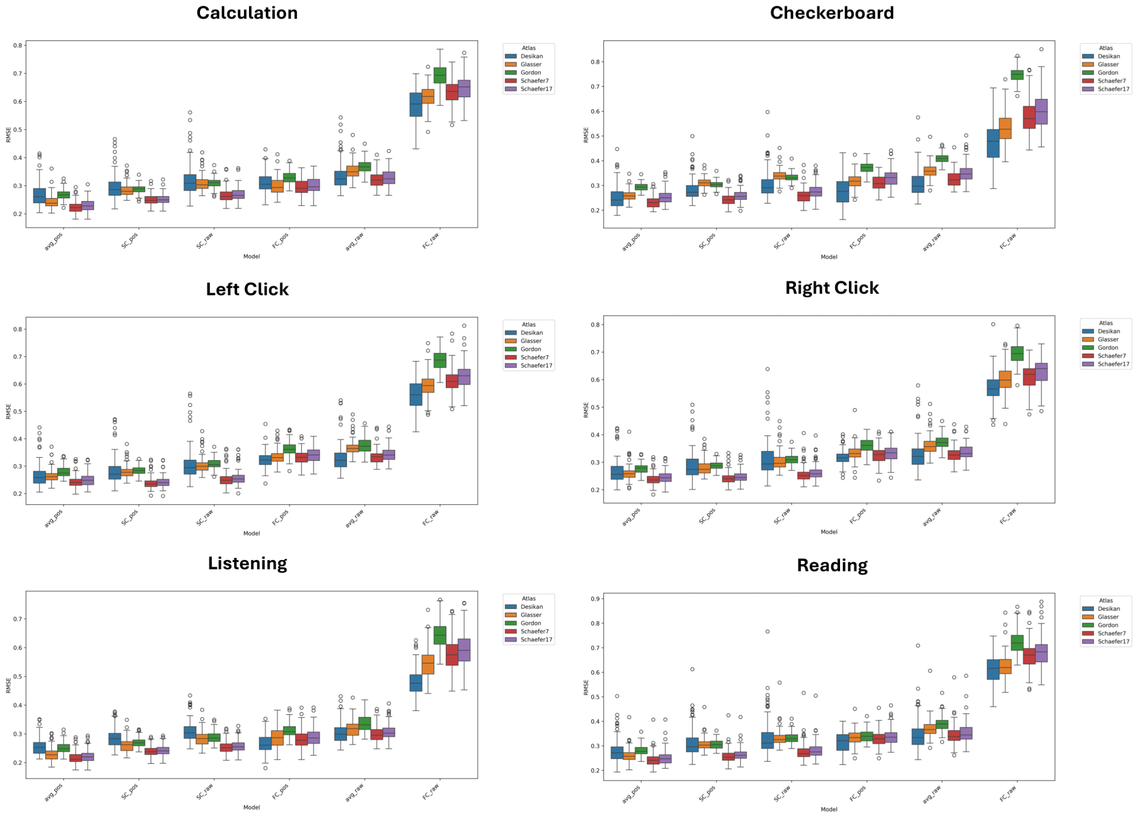


**Figure S6.** Comparison of Mapping Errors (RMSE) Across Multiple Biological Models, Tasks, and Parcellations (Study 1)

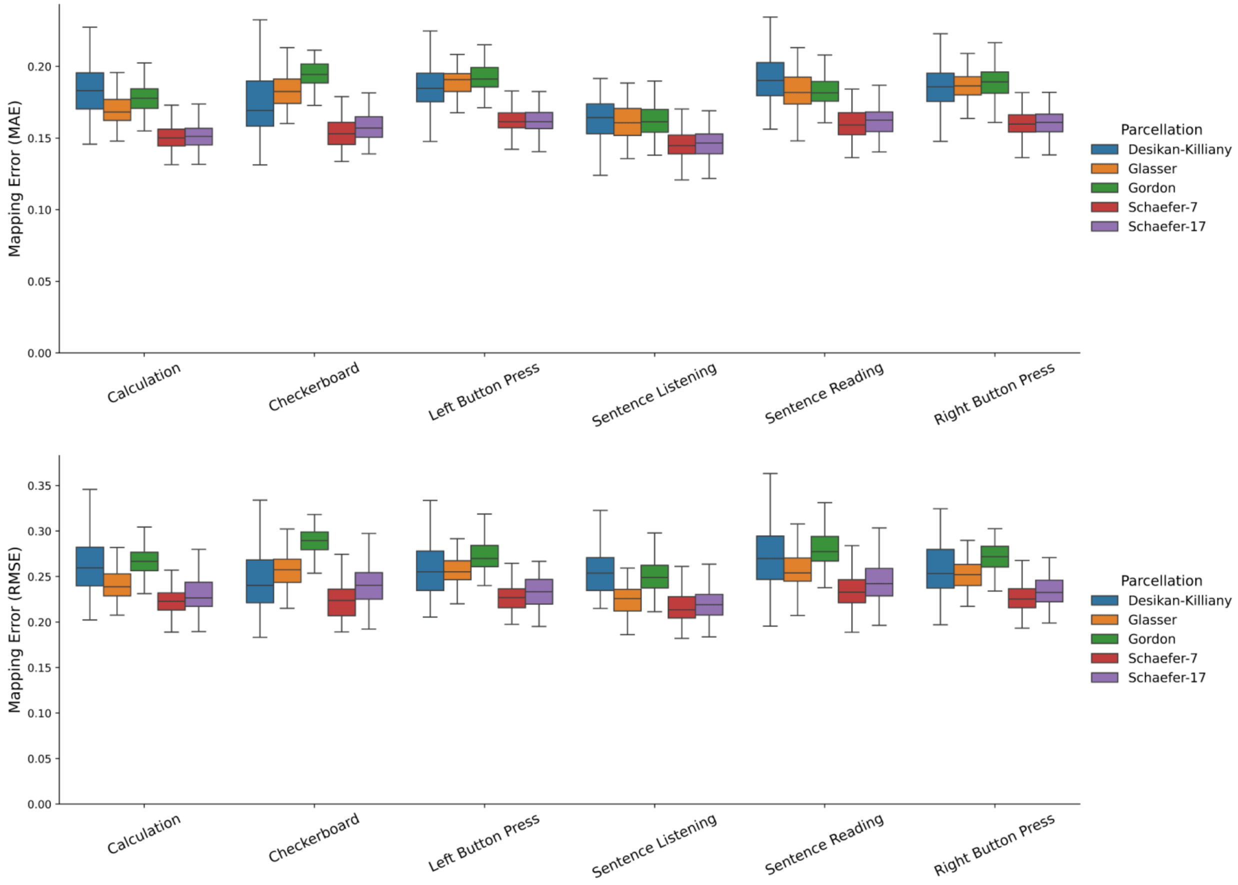


**Figure S7.** Mapping Errors (MAE, RMSE) of the Best Fitting Model for Each Parcellations by Task Conditions (Study 1).

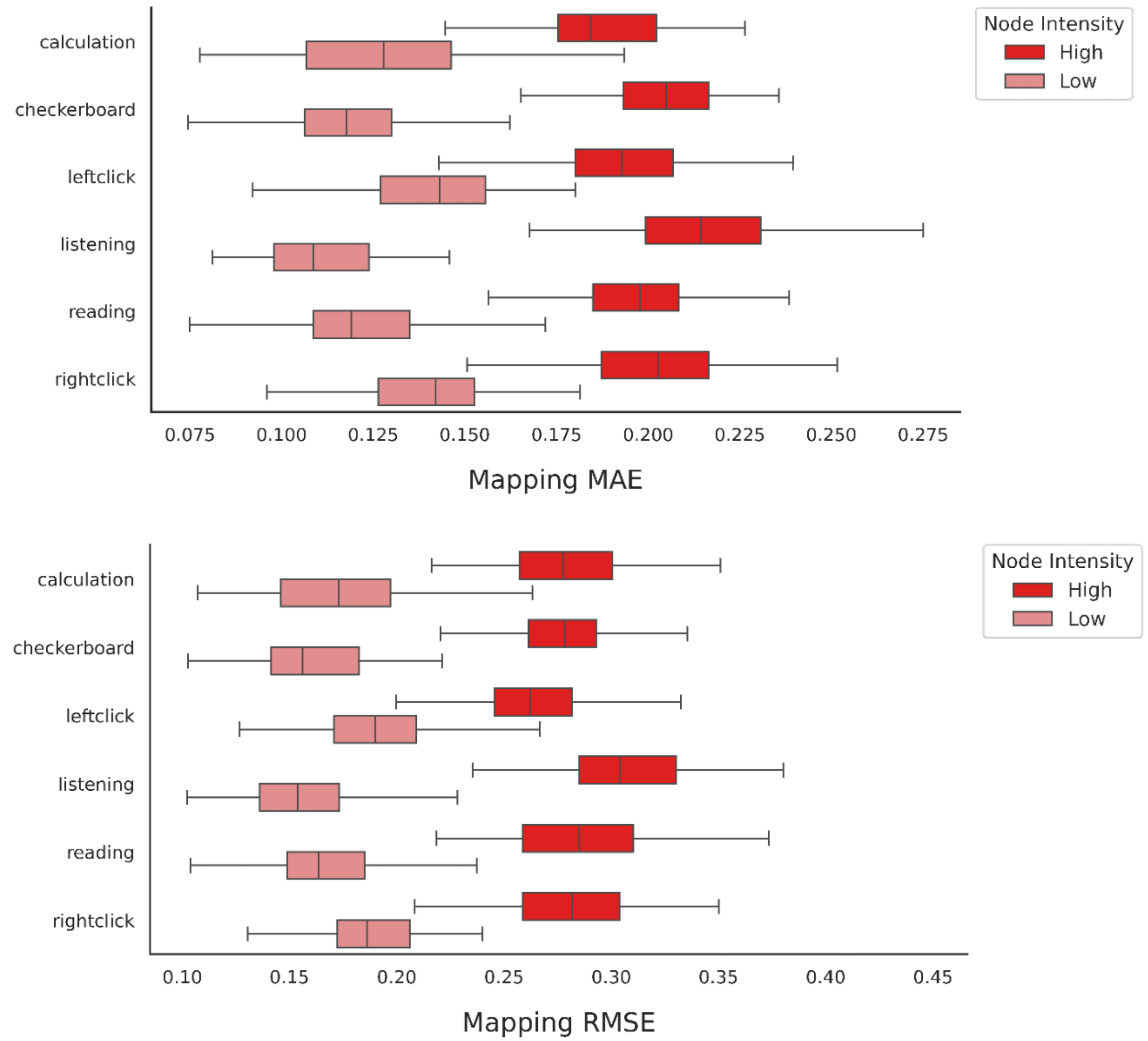


**Figure S8.** Impact of signal intensity on Mapping Errors (MAE, RMSE, Study 1). Bar charts shows differences in mapping errors when analyses were restricted to nodes in the top 25% highest signal intensity and the bottom 25% lowest signal intensity for each task condition.

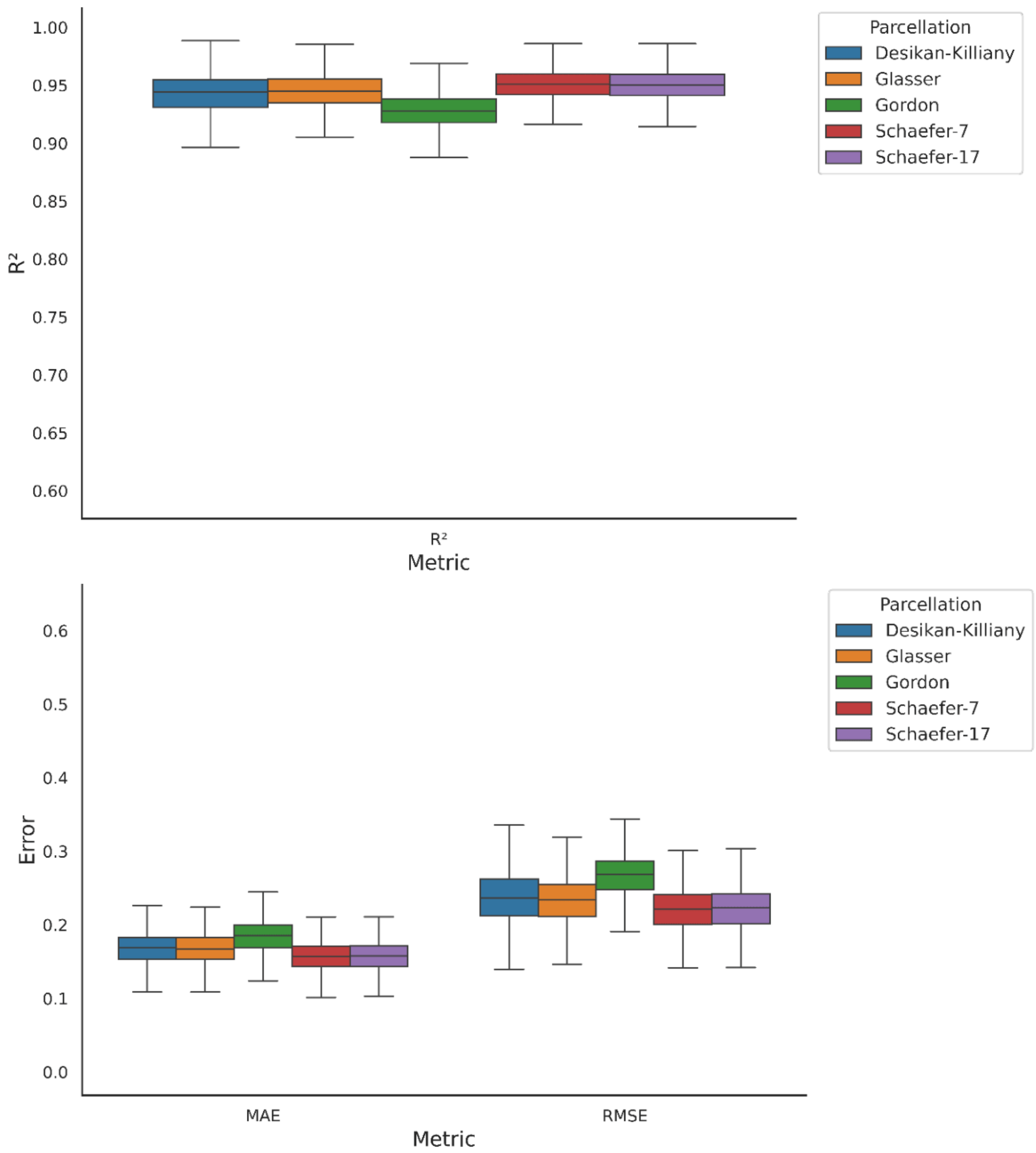


**Figure S9.** Mapping Accuracy ($R^2$) & Errors (MAE, RMSE) of the best fitting propagation model on 1,000 randomly selected functional MRI brain images.

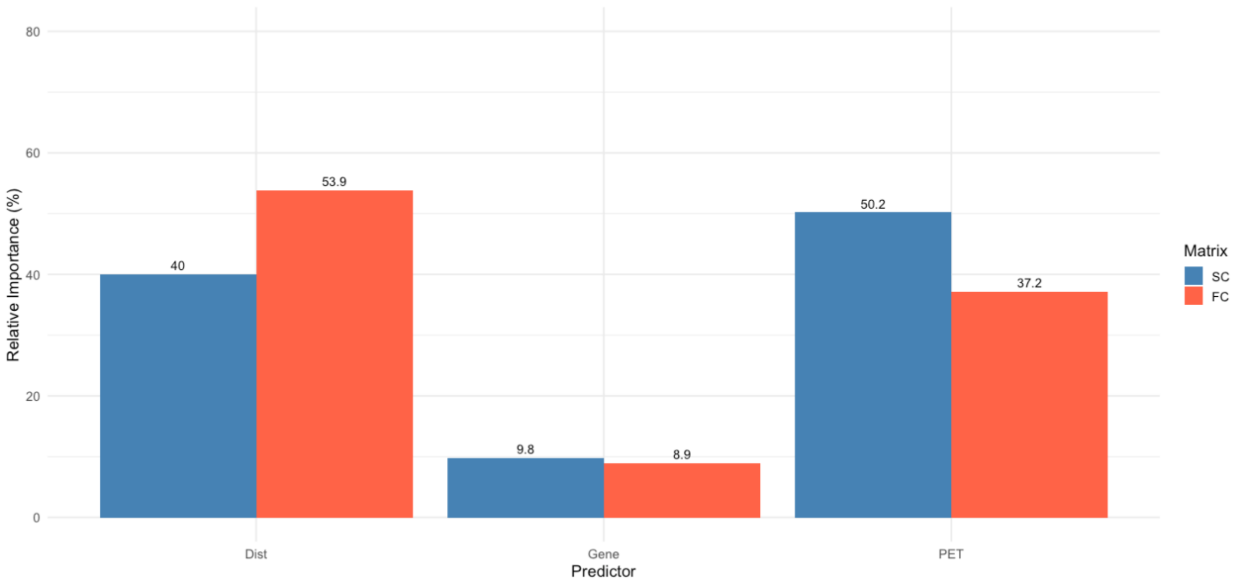


**Figure S10.** Post hoc Dominance Analyses on Contributors to Structural Covariance and Functional Connectivity. Graph shows the relative contributions of (a) spatial proximity (Dist), (b) transcriptomic similarity (Gene), and (c) receptor similarity (PET) between Schaefer-400 7-Network cortical parcels to SC and FC edges.

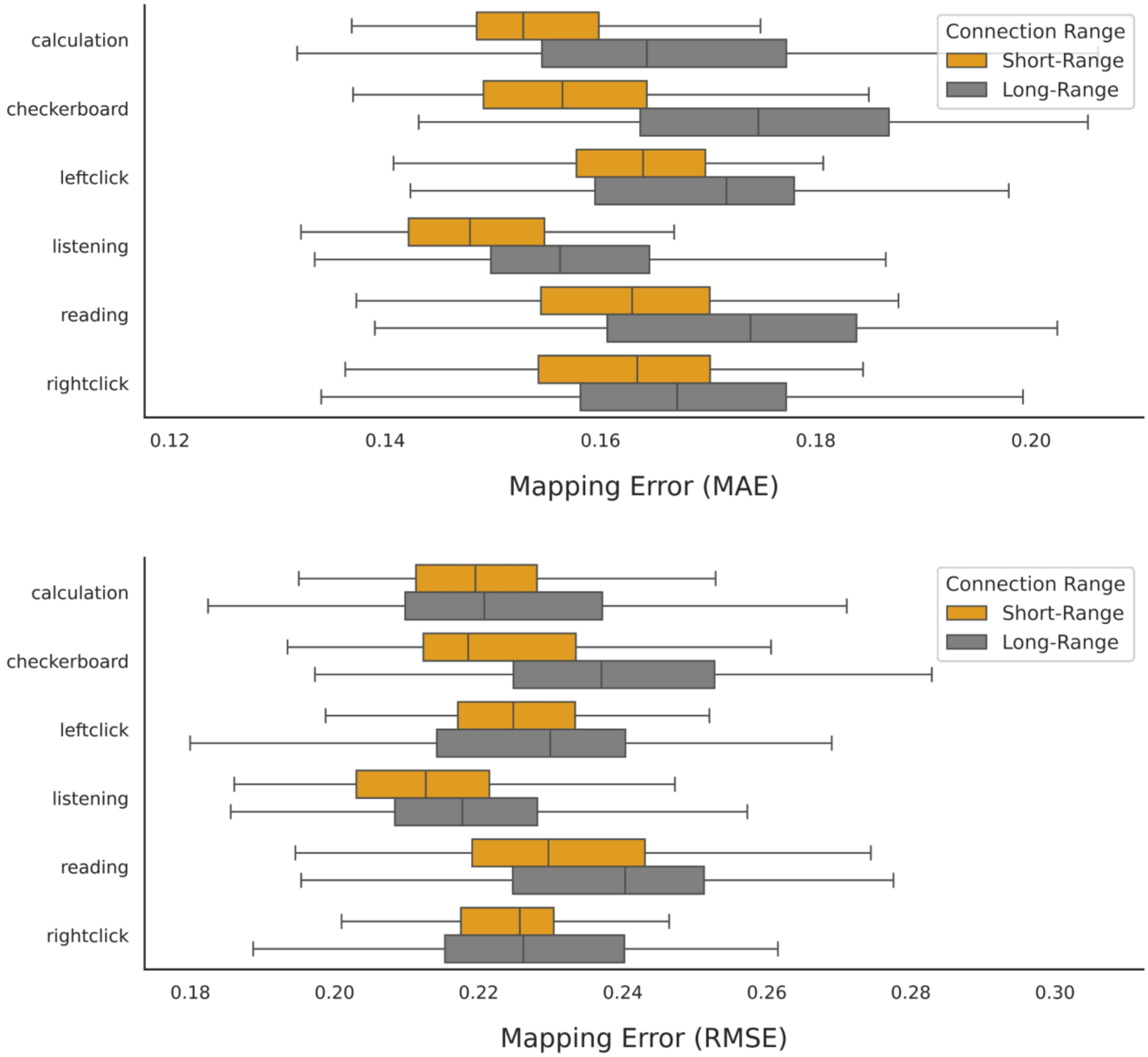


**Figure S11.** Impact of spatial proximity on Mapping Errors (Study 1). To test whether performance was driven by short-range connections, distance-based cross-validation was conducted. The training set consisted of the 75% of regions nearest to each source node in surface space, and the test set consisted of the 25% of regions farthest from the source node. Errors of propagation mapping were aggregated across each source node iterations for the training (short-range connections) and test (long-range connections) sets for each subject.

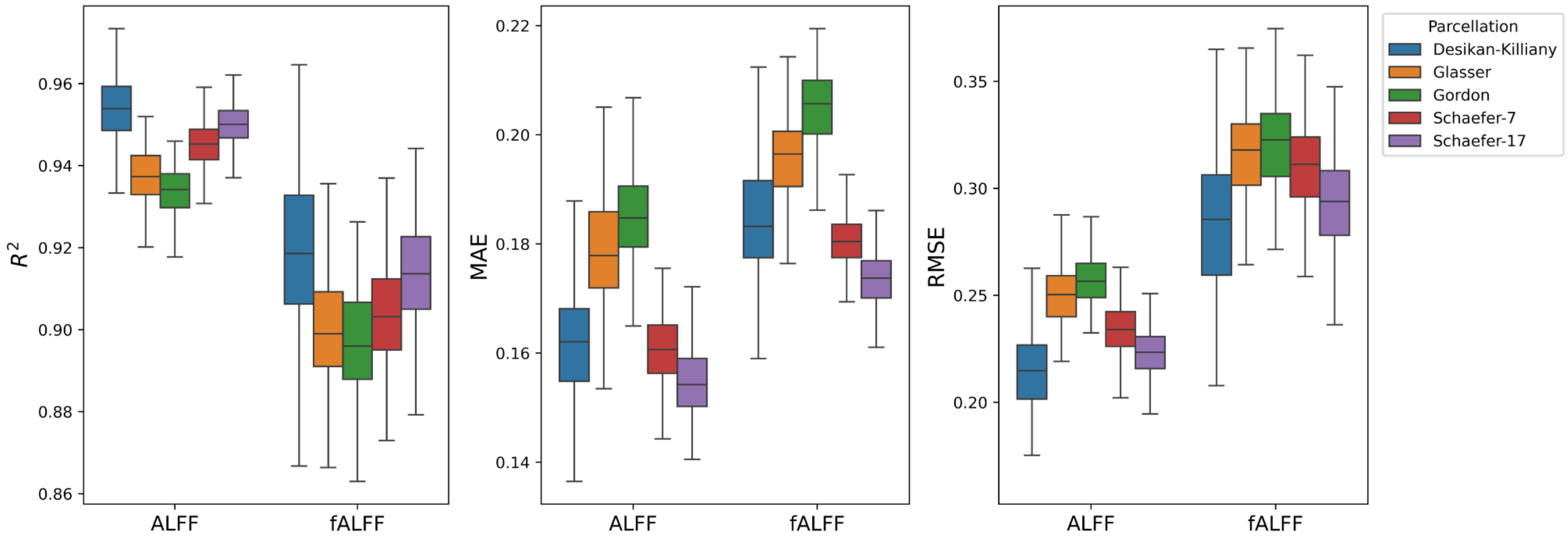


**Figure S12.** Mapping accuracy and errors in an independent sample using the amplitude of low-frequency oscillations during resting-state (Study 2)

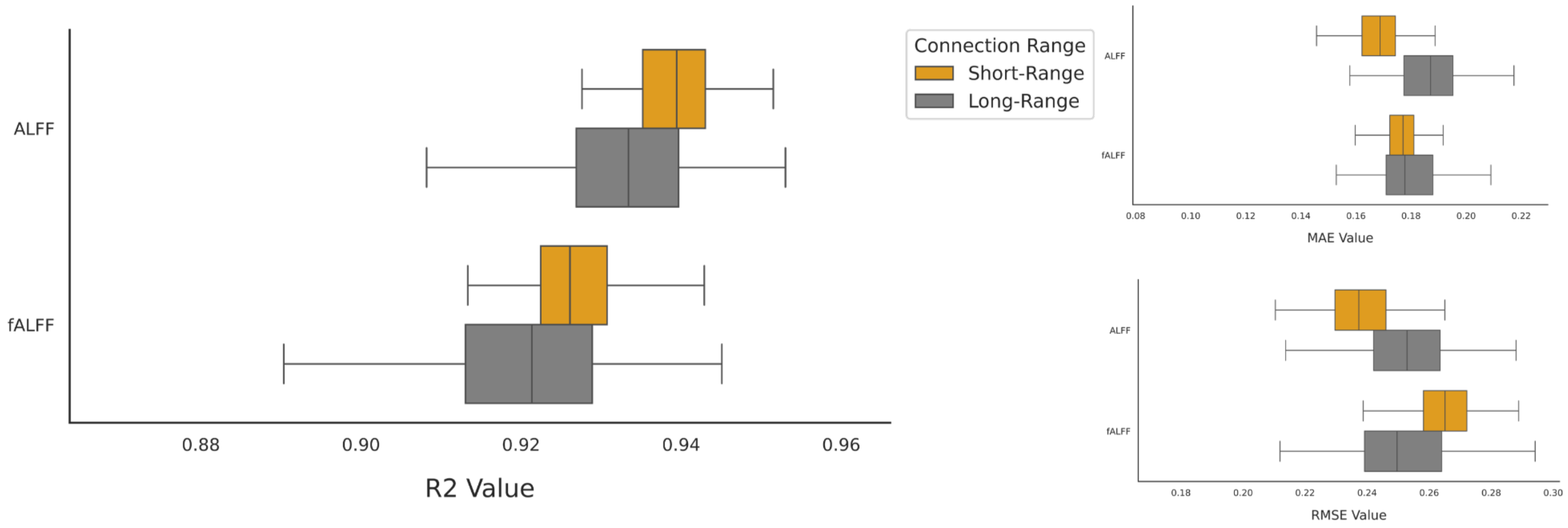


**Figure S13.** Impact of spatial proximity on mapping accuracy and errors (Study 2). To test whether performance was driven by short-range connections, distance-based cross-validation was conducted. The training set consisted of the 75% of regions nearest to each source node in surface space, and the test set consisted of the 25% of regions farthest from the source node. Errors of propagation mapping were aggregated across each source node iterations for the training (short-range connections) and test (long-range connections) sets for each subject.

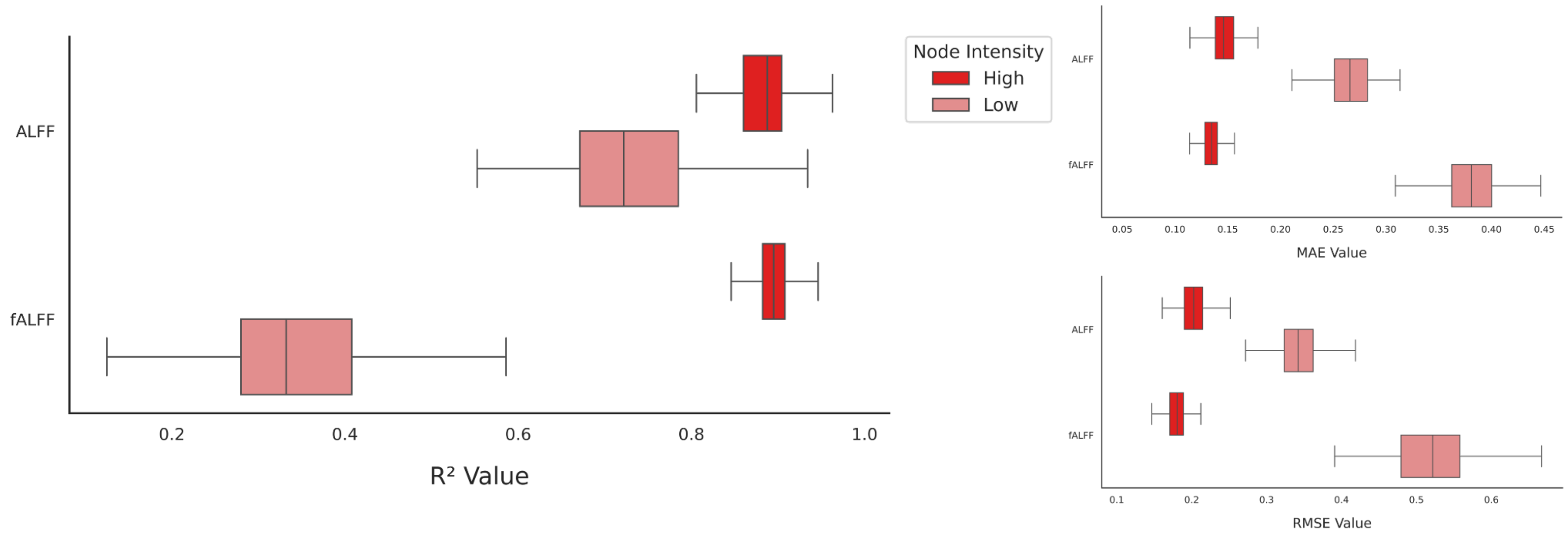


**Figure S14.** Impact of Signal Intensity on Mapping Accuracy and Errors (MAE, RMSE, Study 2). Bar charts shows differences in mapping accuracy and errors when analyses were restricted to nodes in the top 25% highest signal intensity and the bottom 25% lowest signal intensity for each task condition.

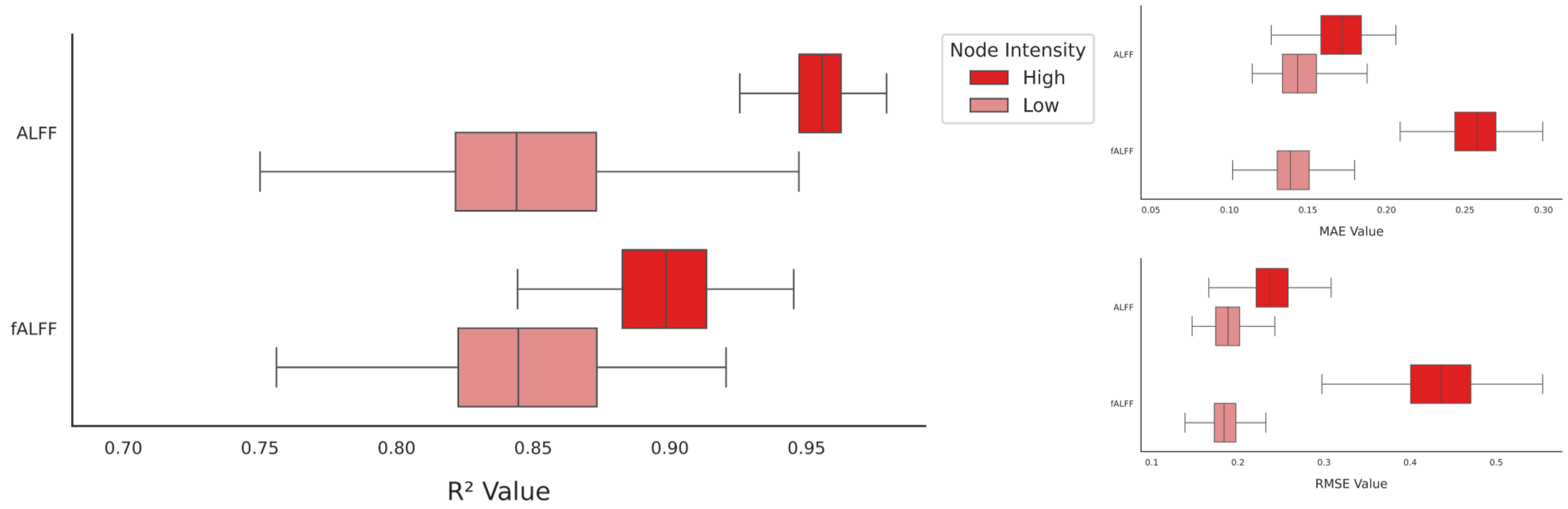


**Figure S15.** Impact of Signal Intensity on Mapping Accuracy and Errors (MAE, RMSE, Study 2). Bar charts shows differences in mapping accuracy and errors when analyses were restricted to nodes in the top 25% highest signal intensity and the bottom 25% lowest signal intensity **in absolute scale** for each task condition.
